# Landscape and Selection of Vaccine Epitopes in SARS-CoV-2

**DOI:** 10.1101/2020.06.04.135004

**Authors:** Christof C. Smith, Sarah Entwistle, Caryn Willis, Steven Vensko, Wolfgang Beck, Jason Garness, Maria Sambade, Eric Routh, Kelly Olsen, Julia Kodysh, Timothy O’Donnell, Carsten Haber, Kirsten Heiss, Volker Stadler, Erik Garrison, Oliver C. Grant, Robert J. Woods, Mark Heise, Benjamin G. Vincent, Alex Rubinsteyn

## Abstract

There is an urgent need for a vaccine with efficacy against SARS-CoV-2. We hypothesize that peptide vaccines containing epitope regions optimized for concurrent B cell, CD4^+^ T cell, and CD8^+^ T cell stimulation would drive both humoral and cellular immunity with high specificity, potentially avoiding undesired effects such as antibody-dependent enhancement (ADE). Additionally, such vaccines can be rapidly manufactured in a distributed manner. In this study, we combine computational prediction of T cell epitopes, recently published B cell epitope mapping studies, and epitope accessibility to select candidate peptide vaccines for SARS-CoV-2. We begin with an exploration of the space of possible T cell epitopes in SARS-CoV-2 with interrogation of predicted HLA-I and HLA-II ligands, overlap between predicted ligands, protein source, as well as concurrent human/murine coverage. Beyond MHC affinity, T cell vaccine candidates were further refined by predicted immunogenicity, viral source protein abundance, sequence conservation, coverage of high frequency HLA alleles and co-localization of CD4^+^ and CD8^+^ T cell epitopes. B cell epitope regions were chosen from linear epitope mapping studies of convalescent patient serum, followed by filtering to select regions with surface accessibility, high sequence conservation, spatial localization near functional domains of the spike glycoprotein, and avoidance of glycosylation sites. From 58 initial candidates, three B cell epitope regions were identified. By combining these B cell and T cell analyses, as well as a manufacturability heuristic, we propose a set of SARS-CoV-2 vaccine peptides for use in subsequent murine studies and clinical trials.

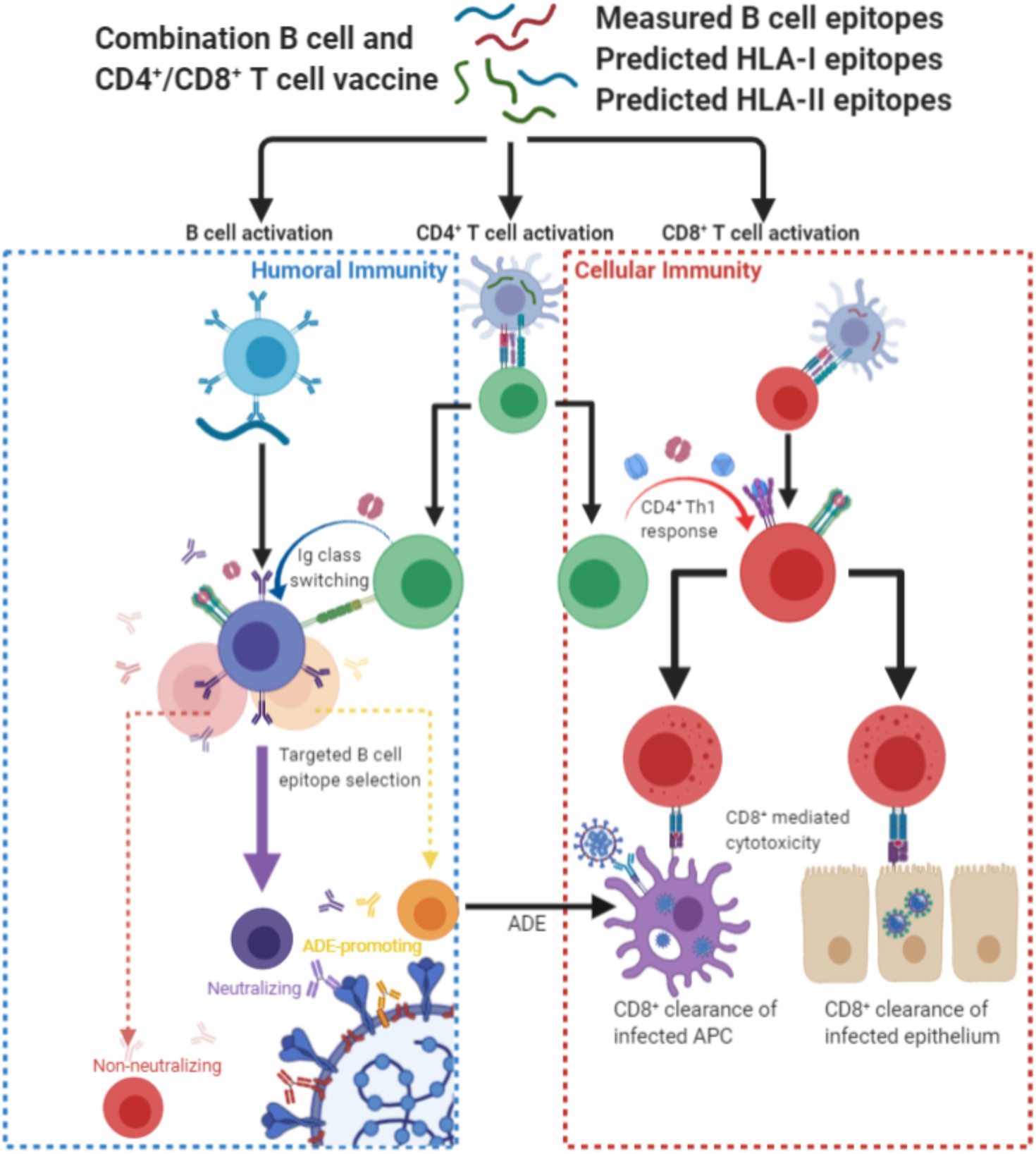

## Introduction

COVID-19, the infectious disease caused by the SARS-CoV-2 virus, is a global pandemic which has infected millions of individuals and caused hundreds of thousands of deaths. Management and treatment options are limited, and development of a vaccine is critical to mitigate public health impact.

SARS-CoV-2 vaccines have largely focused on generation of B cell responses to trigger production of neutralizing antibodies^1–3^. Similar to SARS-CoV-1, SARS-CoV-2 enters cells through interaction of the viral receptor binding domain (RBD) with angiotensin converting enzyme 2 (ACE2) receptors, found on the surface of human nasopharyngeal, lung, and gut mucosa^4^. Production of neutralizing antibodies targeting the RBD or other functional domains is thought to be critical for vaccine efficacy. Generation of non-neutralizing antibody responses may be associated with vaccine failure, and in the worst case scenario enhanced disease upon viral exposure, either through the induction of enhanced pulmonary inflammation^5^, or Fc receptor-mediated antibody-dependent enhancement (ADE)^6^. While anti-SARS-CoV-2 antibodies have been identified in COVID-19 patients, it is unknown which of these antibodies drive viral neutralization, ADE, or both. Thus, vaccine efficacy and safety will be optimized by approaches that maximize generation of neutralizing antibodies while minimizing ADE or pulmonary immune pathology.

In addition to targeting a B cell response, a SARS-CoV-2 vaccine should also drive T-cell activity, because 1) CD4^+^ and CD8^+^ T cells have well-defined roles in the antiviral immune response, including against SARS-CoV-1^7–9^, and 2) CD8^+^ T cells may be able to clear infected antigen presenting cells to mitigate clinical sequelae of ADE or Th2 T cell driven pulmonary immune pathology^5^. Prior studies in SARS-CoV-1 have demonstrated T cell responses against viral epitopes, with strong T cell responses correlated with generation of higher neutralizing antibody titers^9^. Unlike antibody epitopes, T cell epitopes need not be limited to accessible regions of surface proteins. In SARS-CoV-1, concurrent CD4^+^ and CD8^+^ activation and central memory T cell generation were induced in exposed patients; however, increased Th2 cytokine polarization was observed in patients with fatal disease^9^. Thus, vaccines targeting humoral (B cells) and cytotoxic arms (CD8^+^ T cells) with concurrent helper signalling (CD4^+^ T cells), delivered with adjuvants promoting Th1 polarization, may provide optimal immunity against SARS-CoV-2.

Current vaccine strategies in SARS-CoV-2 include recombinant spike (S) glycoprotein, recombinant receptor binding domain (RBD), nucleic acid (DNA and RNA) encodings of the S glycoprotein, adenovirus vector expressing the surface glycoprotein, live recombinant measles vaccine altered to express the surface glycoprotein, as well as delivery of whole inactivated virus^2,3,10–13^. Many of these strategies are attractive for eliciting antibody responses against conformational epitopes. Multi-epitope peptide vaccines are an alternative approach which has a history of safe administration, may be developed and updated rapidly, and may be less likely to elicit non-neutralizing antibodies that contribute to antibody-dependent enhancement (ADE)^14–16^.

We report here a comprehensive survey of the T and B cell epitope space of SARS-CoV-2 (**Figure 1**). Predicted T cell epitopes were derived from *in silico* predictions filtered on binding affinity and immunogenicity models generated from epitopes deposited in the Immune Epitope Database (IEDB)^17^, population diversity, and source protein abundance. B cell epitope candidates were curated from linear epitope mapping studies and further filtered by accessibility, glycosylation, polymorphism, and adjacency to functional domains. Given the rapid development of murine-adapted SARS-CoV-2 models, we also report T cell epitopes predicted to bind murine MHC coded for by H2-D/K^b/d^ and H2-IA^b/d^ haplotypes. We have integrated these data and present a strategy for epitope prioritization for vaccine development.

**Figure 1:**
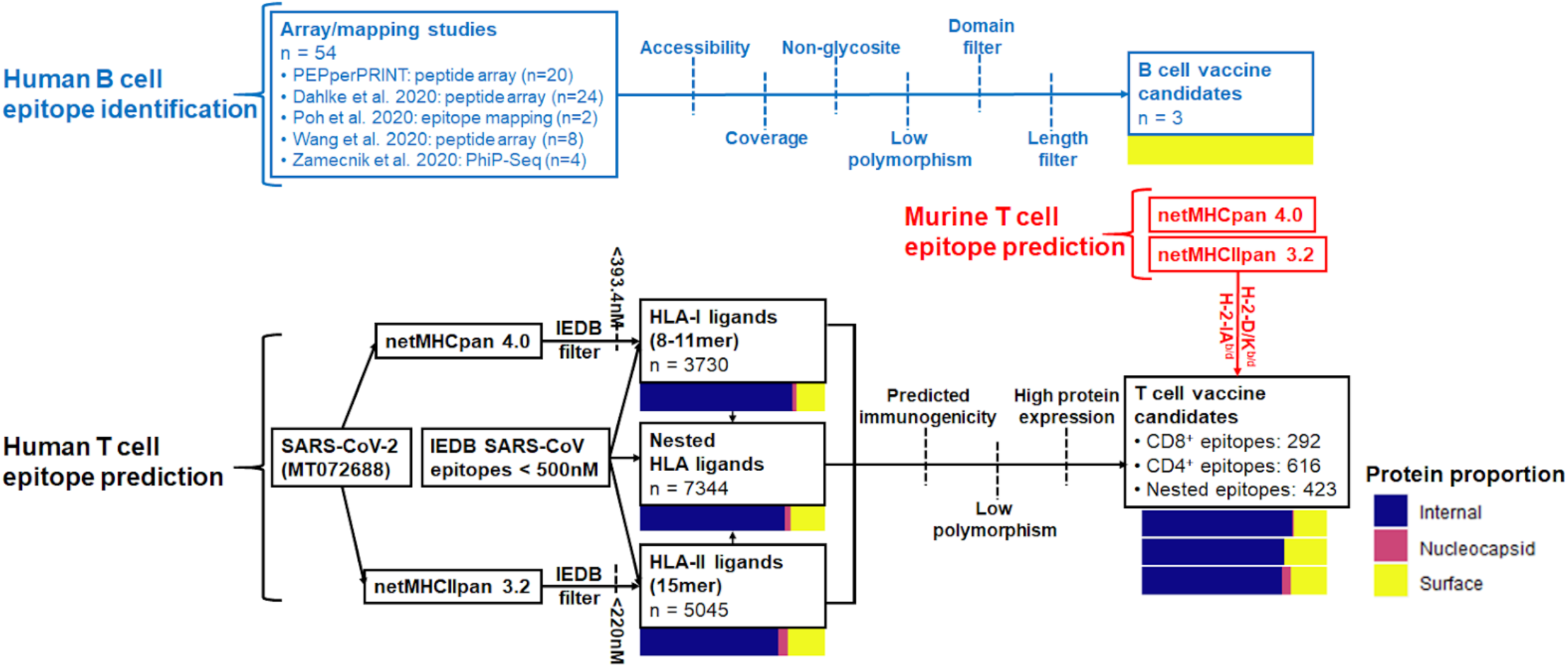
Summary of B cell and CD4^+^/CD8^+^ epitope prediction workflows. Pathways are colored by B cell (blue), human T cell (black), and murine T cell (red) epitope prediction workflows. Color bars represent proportions of epitopes derived from internal proteins (ORF), nucleocapsid phosphoprotein, and surface-exposed proteins (spike, membrane, envelope).

## Results

### Landscape of MHC ligands in SARS-CoV-2

To determine the landscape of potential HLA ligands in SARS-CoV-2 (**Figure 1**, black), we first identified candidate MHC ligands by performing HLA-I binding prediction using NetMHCpan 4.0 (both EL and BA mode)^18^ and MHCflurry^19^ (8-11mers), and HLA-II binding prediction using NetMHCIIpan 3.2^20^ and 4.0^21^ (15mers), using alleles with >5% genetic frequency in the United States^22,23^ (full predicted sets: **Table S1, S2**). To assess the accuracy of these peptide/MHC binding prediction tools on viral peptides, we tested their performance on IEDB MHC affinity assay data values for viral peptides. Of the predictive models evaluated, NetMHCpan 4.0 (BA) and NetMHCIIpan 3.2 demonstrated the highest correlation of binding affinity predictions for Class I and Class II MHC, respectively (**Figure S1A-B**). Therefore, these two predictors were for predicting MHC ligands. A measured peptide/MHC binding affinity of 500 nM or less is commonly used to identify MHC-binding peptides which are more likely to be T cell epitopes^24^. To account for the inaccuracy inherent to prediction (as opposed to measurement) of peptide-MHC affinity, we derived slightly stricter cutoffs. In order to achieve 90% specificity in IEDB binding affinity data, we use predicted binding affinity thresholds of 393.4 nM and 220.0 nM for Class I and Class II MHC, respectively, (**Supplementary Figure 1C-D**). This filter was applied to NetMHCpan 4.0 and NetMHCIIpan 3.2 SARS-CoV-2 MHC binding predictions, which removed the majority of viral protein sub-sequences (**Figure 2A-B**).

**Figure 2:**
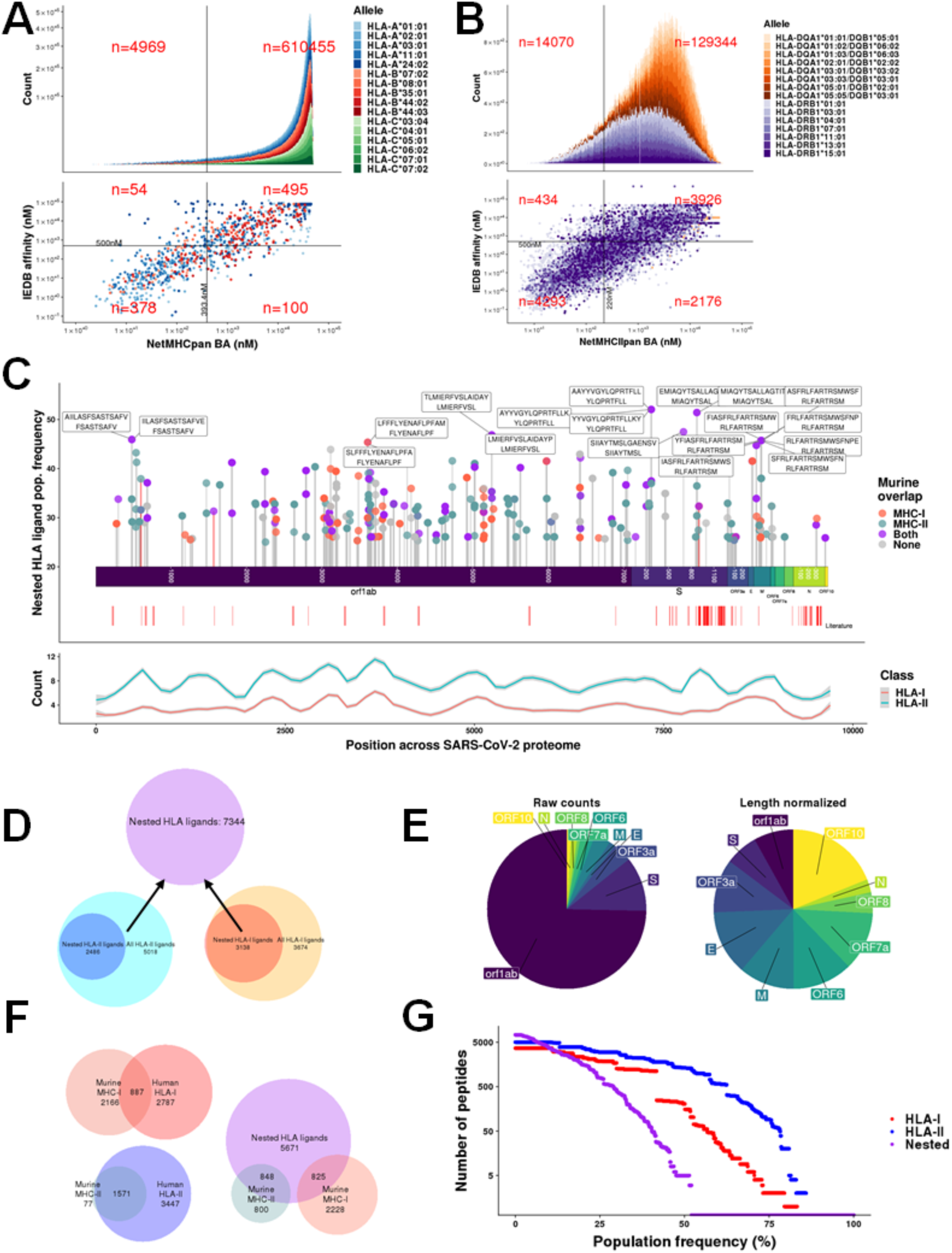
Landscape of SARS-CoV-2 MHC ligands. **(A&B)** Selection criteria for **(A)** HLA-I and **(B)** HLA-II SARS-CoV-2 HLA ligand candidates. Scatterplot (bottom) shows predicted (x-axis) versus IEDB (y-axis) binding affinity, with horizontal line representing 500nM IEDB binding affinity and vertical line representing corresponding predicted binding affinity for 90% specificity in binding prediction. Histogram (top) shows all predicted SARS-CoV-2 HLA ligand candidates. **(C)** Landscape of predicted HLA ligands, showing nested HLA ligands comprising HLA-I and -II ligands with complete overlap (top), and LOESS fitted curve (span = 0.1) for HLA-I/II ligands by location along the SARS-CoV2 proteome (bottom). Red track represents SARS epitopes identified in literature review with sequence identity in SARS-CoV-2. Predicted HLA ligands with conserved sequences to this literature set are represented in the lollipop plot with a red stick. **(D)** Summary of total number of predicted HLA-I/II ligands and nested HLA ligands. **(E)** Summary of nested HLA ligand coverage by protein, with raw counts (left) or counts normalized by protein length (right). **(F)** Summary of murine/human MHC ligand overlap. **(G)** Distribution of population frequencies among predicted HLA-I, -II, and nested HLA ligands.

With the goal of finding epitope regions capable of inducing both CD4^+^ and CD8^+^ T cell responses, we analyzed our MHC ligand predictions for the set of overlapping HLA-I and HLA-II ligand combinations, referred to here as nested HLA ligands. To generate these nested HLA ligands, each predicted HLA-I ligand was paired with an HLA-II ligand with full sequence overlap, selecting for the HLA-II ligand(s) with highest population coverage (**Figure 2C,D**, n = 7344 pairs consisting of 2486 unique HLA-I ligand and 3138 unique HLA-II ligands). Predicted MHC ligands were not evenly distributed across the proteome, with local peaks and troughs observed that correlated between HLA-I and -II ligands (**Figure 2C**, bottom; Pearson correlation of HLA-I/II LOESS (span = 0.1): r = 0.703, p < 0.001). Notably, while SARS-CoV-1 T cell epitopes previously described in the literature were primarily located in the surface glycoprotein (S) and nucleocapsid protein (N) (**Table S3**)^9,25–50^, we observed a paucity of predicted MHC ligands in the nucleocapsid protein (N). Of 113 unique T cell epitopes described in the literature that were also found in the SARS-CoV-2 proteome, we observed only two HLA-I peptide sequences in our predicted nested HLA ligand set. Numbers of predicted nested MHC ligands were associated with protein length (**Figure 2E**, left), with orf1ab having the greatest count; however, normalizing by protein length demonstrated greater equality of distribution, with the three largest viral proteins (orf1ab, S, and N) being among the lowest ranked (**Figure 2E**, right).

As murine models for SARS-CoV-2 would be a powerful tool in understanding viral immunobiology, we determined which predicted HLA ligands were also predicted to bind murine H2-b/d MHC. NetMHCpan and NetMHCIIpan were run using the SARS-CoV-2 proteome against the H2-b and H2-d haplotypes, filtering by MHC-I ligands top 2nd percentile (n = 3053) and MHC-II ligands in the top 10th percentile (n = 1648). From this set, we observed an overlap of 887 peptides in MHC-I and 1571 peptides in MHC-II between murine and human sets (**Figure 2F**). For the nested HLA ligand set, we observed 825 and 848 overlapping murine MHC-I and -II ligands, respectively, with 846 HLA ligands containing both murine MHC-I and -II coverage.

The majority of HLA ligand sequences were predicted to bind to fewer than 50% of the U.S. population, particularly for HLA-I and nested ligands (**Figure 2G**). In accordance with higher population coverage distribution in HLA-II, predicted HLA-II ligands also demonstrated more binding alleles on average (mean alleles per peptide: HLA-I = 1.35, HLA-II = 2.80). Among the most common alleles were HLA-A*02:01 (n = 784), HLA-A*11:01 (n = 643), and HLA-A*03:01 (n = 383) for predicted HLA-I binding peptides and HLA-DRB1*01:01 (n = 5401), HLA-DRB1*07:01 (n = 3225), and HLA-DRB1*13:01 (n = 3022) for predicted HLA-II binding peptides.

### CD8^+^ and CD4^+^ T cell epitope prediction

Peptide/MHC binding is necessary but not sufficient for peptide epitopes to elicit T cell responses. We sought to identify a set of epitopes that would serve as good targets for a SARS-CoV-2 T cell vaccine. From the total pool of HLA-I, HLA-II, and nested MHC ligands, we sought to prioritize sequences which are predicted to be immunogenic from highly conserved regions of abundant viral proteins (**Figure 3 middle**).

**Figure 3:**
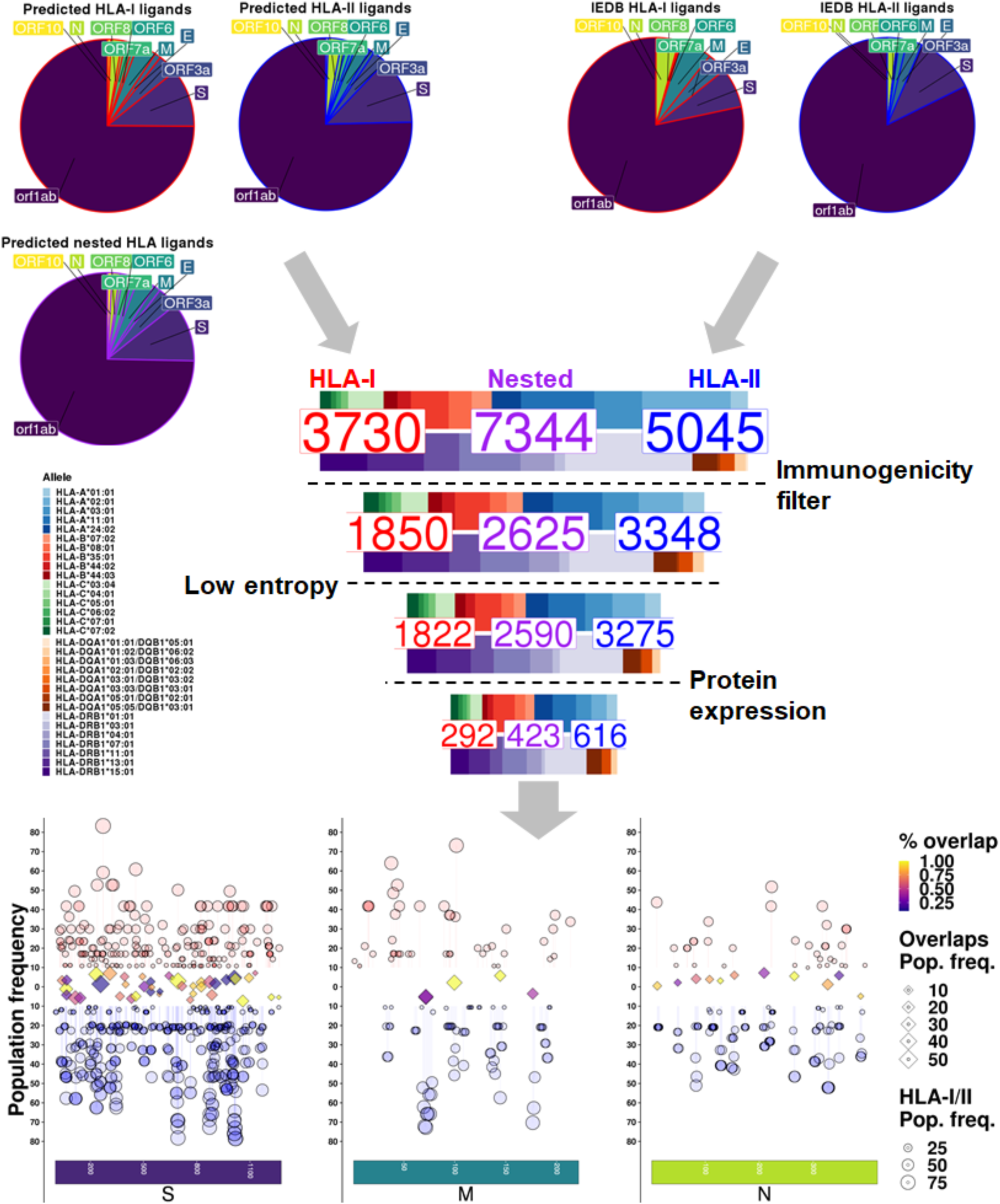
Prediction of SARS-CoV-2 T cell epitopes. **(Top)** Summary of predicted (left) and IEDB-defined (right) SARS-CoV-2 HLA ligands, showing proportions of each derivative protein. **(Middle)** Funnel plot representing counts of HLA-I (red text), HLA-II (blue text), and nested HLA (violet text) ligands along with proportions of HLA-I (top bar) and HLA-II (bottom bar) alleles at each filtering step. **(Bottom)** Summary of CD8^+^ (red, top), CD4^+^ (blue, bottom), and nested T cell epitopes (middle) after filtering criteria in S, M, and N proteins. Y-axis and size represent the population frequency of each CD8^+^ and CD4^+^ epitopes by circles. Middle track of diamonds represents overlaps between CD8^+^ and CD4^+^ epitopes, showing the overlap with greatest population frequency (size) for each region of overlap. Color of diamonds represents the proportion of overlap between CD4^+^ and CD8^+^ epitope sequences.

To predict the immunogenicity of MHC ligands, we fit a forward stepwise multivariable logistic regression model using peptide/HLA tetramer flow cytometry data curated from viral entries of the Immune Epitope Database (IEDB)^17^. Tetramer data was selected for the response variable because it provides unambiguous association between a peptide and its bound MHC, and additionally tests which specific peptide/MHC is capable of eliciting a T cell response. Each unique peptide-MHC was encoded with features derived from epitope prediction tools as well as features relating to amino acid content (See *Methods: Immunogenicity modeling*). Model performance in 5-fold cross validation demonstrated AUC values of approximately 0.7 and 0.9 for HLA-I and -II, respectively, in both training and test sets (**Figure S2A-B**). Models demonstrated cleaner separation of tetramer positive and negative groups for CD4^+^ epitopes compared to HLA-I (**Figure S2C-D**). To determine a cause for this difference in model performance, we examined predicted binding affinity scores between tetramer positive and negative epitopes, which demonstrated significantly better separation for CD4^+^ epitopes than CD8^+^ epitopes (**Figure S2E-F**). In accordance with this difference in binding affinity distribution, the HLA-II model showed strong association between lower binding affinity and lower predicted tetramer positivity, while the HLA-I model showed a weaker inverse association (**Figure S3**). Due to these binding affinity distribution differences between IEDB HLA-I and HLA-II tetramer sets, a performance-based cutoff did not allow for equal filtering of CD4^+^ and CD8^+^ epitopes. Therefore, we filtered by GLM scores above the median in each HLA-I/II SARS-CoV-2 epitope group, which provided balanced selection while removing predicted low-immunogenicity epitopes (**Figure S4**).

Next, we sought to prioritize epitopes derived from regions of low sequence variation across viral strains. A position-based entropy filter was applied to all epitopes (**Figure S5**), keeping those with an entropy score ≤ 0.1 (approximately 98% sequence identity) in all amino acid positions across MSA-aligned SARS-CoV-2 genomes within the Nextstrain database^51,52^. High entropy was observed in the well-described spike protein D614G polymorphic site (**Figure S5A**, red dot). Other areas of high entropy included positions 3606, 4715, 5828, and 5865 of orf1ab, and position 84 of ORF8 (all with entropy > 0.4). The majority of positions demonstrated >95% sequence identity, suggesting high homology between different SARS-CoV-2 viral genomes (**Figure S4B**). Lastly, as the likelihood of MHC presentation is correlated with protein expression^53^, we filtered epitopes to those derived from the three highest expressed SARS-CoV-2 proteins normalized by protein length (**Figure S6**)^54^. Protein abundance was determined from both semi-quantitative mass spectrometry and RNA-seq data^54,55^. After all these filtering steps, 292 CD8^+^, 616 CD4^+^ and 423 nested T cell epitopes were predicted. Relative proportions of HLA alleles were conserved throughout filtering (**Figure 3, middle**). Full peptide sets with all filtering criteria are listed in **Tables S1** (HLA-I) and **S2** (HLA-II).

### B cell epitope prediction

In addition to identifying SARS-CoV-2 T cell epitopes, we sought to identify a set of linear B cell epitopes on the Spike protein which would serve as good targets for stimulating neutralizing antibody responses (**Figure 1**). Epitope candidates were derived from four published preprint mapping/array studies^56–59^ and one as-of-yet unpublished PEPperCHIP® peptide array study (for study details see *Methods: Antibody epitope curation*). Starting with an initial candidate pool of 58 linear epitopes with data to support *in vivo* generation in humans (**Figure 4A, Table S4**), we applied a set of filtering criteria to narrow our target space (**Figure 4B**):

**Figure 4:**
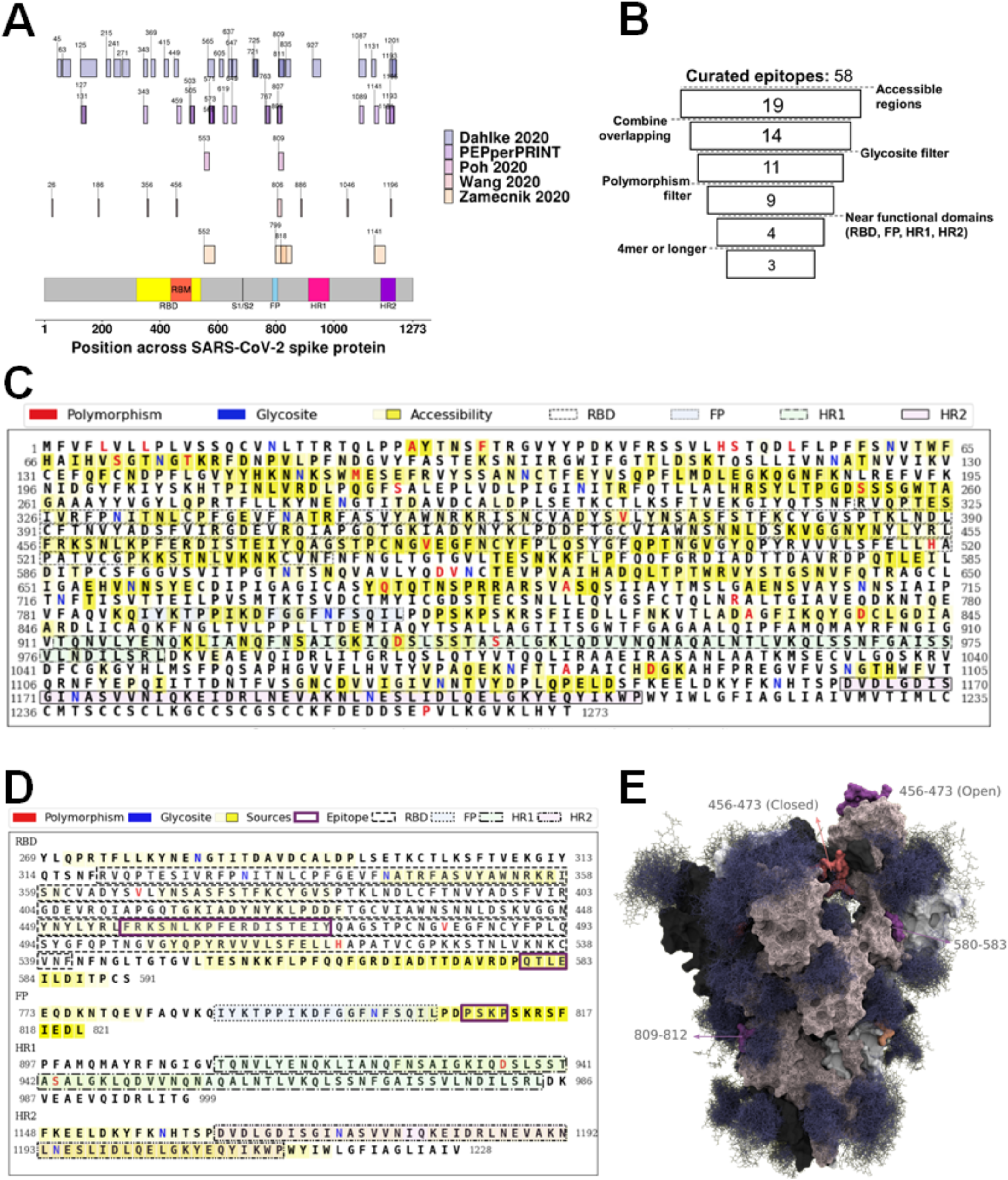
Selection of SARS-CoV-2 B cell epitope regions. **(A)** SARS-CoV-2 linear B cell epitopes curated from epitope mapping studies. X-axis represents amino acid position along the SARS-CoV-2 spike protein, with labeled start sites. **(B)** Schematic for filtering criteria of B cell epitope candidates. **(C)** Spike protein amino acid sequence, with overlay of selection features prior to filtering. Polymorphic residues are red, glycosites are blue, accessible regions highlighted in yellow. The receptor binding domain (RBD), fusion peptide (FP), and HR1/HR2 regions are outlined. **(D)** Spike protein functional regions (RBD, FP, HR1/2) amino acid sequences, with residues colored by how many times they occur in identified epitopes. Selected accessible sub-sequences of known antibody epitopes highlighted in purple outline. **(E)** S protein trimer crystal structure with glycosylation, with final linear epitope regions highlighted by color.

1. Contiguous subsequences of the spike protein with high accessibility
2. Exclude glycosylation sites
3. Exclude regions with significant polymorphism between SARS-CoV-2 strains
4. Keep candidate epitopes within or adjacent to functional domains with evidence of antibody-mediated viral neutralization (RBD, FP, HR1, and HR2)
5. Exclude any candidates shorter than four amino acids

We used SARS-CoV-2 S protein accessibility data from Grant et. al^60^, which resulted in 19 remaining regions after filtering for contiguous stretches with mean accessibility of 35%, minimum accessibility of 15%, requiring at least one residue to have accessibility greater than 50%, and the ends of a region to have at least 25% accessibility. Since many epitopes occur in multiple sources, we combined overlapping epitope candidates into 14 unique sequences. After filtering out epitopes containing glycosites, which may alter antibody binding characteristics^61,62^, 11 non-glycosylated regions remained. Two additional regions were removed because they contained polymorphic sites, defined by mutation frequency > 0.1% from Nextstain SARS-CoV-2 viral sequences. Of the remaining 9 regions, only 4 were close to functional domains which in the closely related virus SARS have evidence of antibody-mediated viral neutralization: the receptor binding domain (RBD), fusion protein (FP), and heptad repeat 1 and 2 (HR1/HR2)^63–68^. Adjacency to a functional region was defined as within 15 aa of either side of FP, HR1, and HR2, and within 50 aa of the RBD. A broader window was used for the receptor binding domain due to the known presence of neutralizing antibody epitopes in S1 of SARS outside of the RBD ^69^. This filtering resulted in four remaining regions, of which our final criteria removed one which had length less than four residues (**Figure 4B**). This filtering criteria precluded the vast majority of total spike protein regions (**Figure 4C**), with three predicted antibody binding regions (residue lengths 18, 4, and 4) remaining (**Figure 4D**). All three epitope candidate regions were present on solvent-exposed surfaces of the S protein trimer 3D structure (**Figure 4E**). It is worth noting that the largest region, residues 456-473 within the receptor binding motif (RBM) loop is only accessible when the RBD is in the “open” conformation.

### Selection of human and murine SARS-CoV-2 vaccine peptides

With the above filters applied to predicted T and B cell epitope candidates, we sought to derive a collection of minimal recommended peptide sets for all combinations of the following vaccine criteria: optimization for CD4^+^ responses, CD8^+^ responses, and coverage of predicted B cell epitopes. We derived 27mer sequences for these vaccine peptide sets, determining peptide combinations which maximized population coverage of T cell epitopes, with or without additional coverage for murine H2-b, H2-d, or both haplotypes (**Figure 5A-B**). If population coverage was identical for multiple candidates, peptides were also optimized based on a manufacturability difficulty scoring system (**Figure S7**). Optimizing for CD4^+^ epitope population coverage demonstrated 88.5% population frequency encompassed by three 27mer peptides (**Figure 5B:** 1, 9, and 15), while CD8^+^ epitope optimization provided 95.8% population frequency coverage by three 27mer peptides (**Figure 5B:** 1, 4, and 14). CD4^+^/CD8^+^ co-optimization provided the best overall population coverage at 81.6% population frequency with four 27mer peptides (**Figure 5B:** 1, 6, 9, 13). While B cell epitope optimization provided CD8^+^ coverage above 85%, CD4^+^ coverage was only 52.8%, suggesting the design of a combination B cell/CD4^+^ T cell vaccine requires use of non-spatially overlapping sequences. Overall, selection of peptides which also provided both H2-b and H2-d epitope coverage did not greatly impact population coverage, suggesting these murine-encompassing sets may allow for vaccine studies in animal models whilst preserving human relevance. Across the different selection criteria for minimal vaccine peptide sets there was significant redundancy. Collapsing the set of vaccine peptides by unique sequences results in a final set of 22 27mer vaccine peptides (**Figure 5B**). In addition to 27mer peptides, all individual T/B cell epitopes (S, M, and N: **Table S5**; all proteins: **Table S6**) as well as 15mer (**Figure S8**) and 21mer (**Figure S9**) optimized peptide sets are also available.

**Figure 5:**
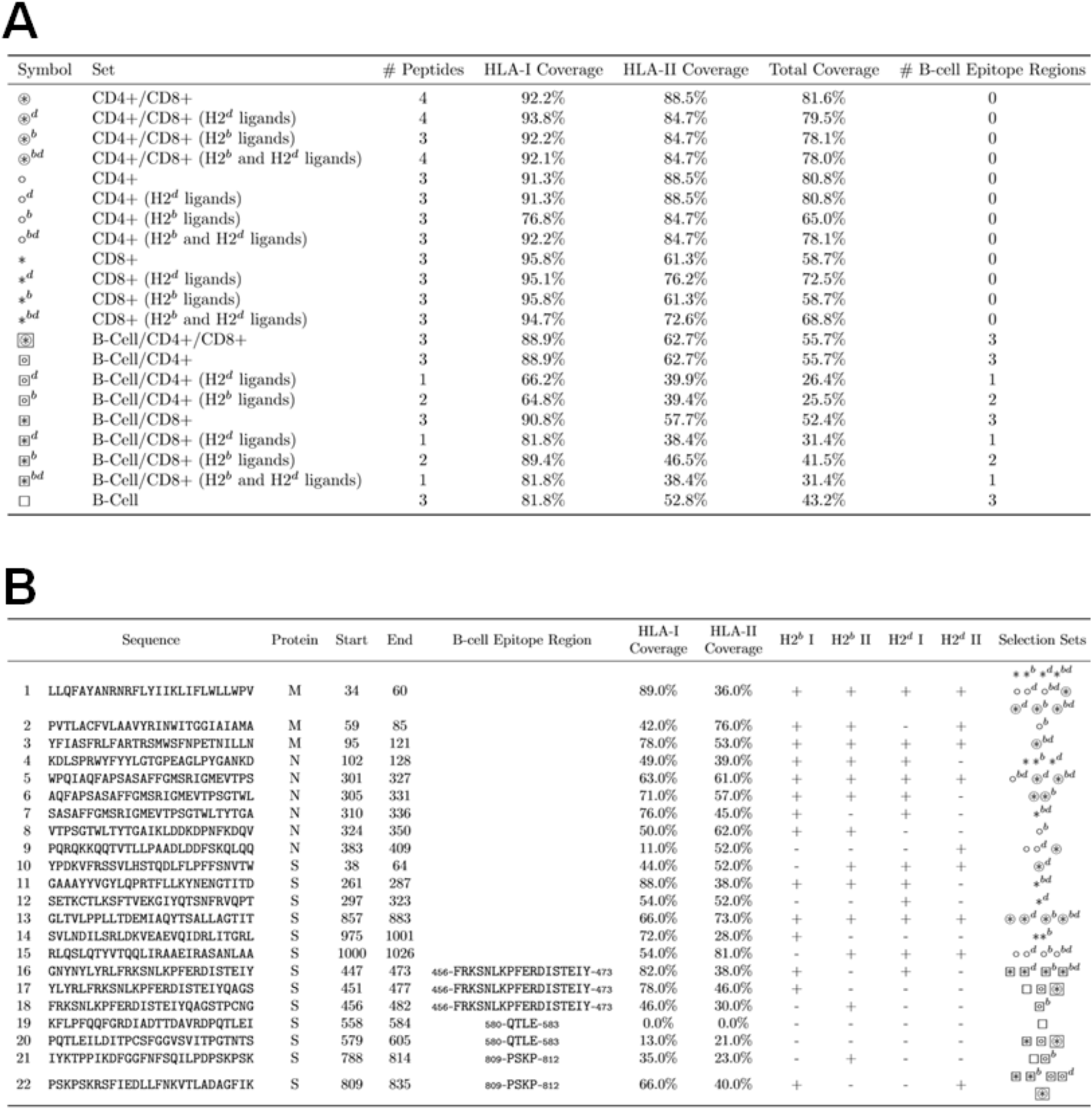
T cell and B cell vaccine candidates. **(A)** 27mer vaccine peptide sets selecting for best CD4^+^, CD8^+^, CD4^+^/CD8^+^, and B cell epitopes with HLA-I, HLA-II, and total population coverage. **(B)** Unified list of all selected 27mer vaccine peptides. Vaccine peptides containing predicted ligands for murine MHC alleles (H2-b and H2-d haplotypes) are indicated in their respective columns.

## Discussion

We report here a survey of the SARS-CoV-2 epitope landscape along with a strategy for prioritizing both T cell and B cell epitopes for vaccine development. Major vaccine efforts targeting coronaviruses have focused on generation of neutralizing antibody responses^70–78^. This is likely critical for vaccine efficacy; however, vaccines against SARS-CoV-2 may also generate non-neutralizing antibodies that facilitate viral entry into cells that express Fc receptor, a phenomenon known as antibody-dependent enhancement (ADE). It is possible that vaccines that elicit a vigorous cytotoxic T cell response will drive early killing of infected cells and mitigate the toxicity associated with ADE. In addition, CD4^+^ T cells provide help to B cells to support class switching, maturation, and antibody production, as well as promoting CD8^+^ T cell activation, maturation, and effector function. In light of this, we searched for vaccine peptide sequences which include both B cell epitopes as well MHC ligands predicted to drive CD4^+^ and CD8^+^ T cell responses at high population frequencies. Our current efforts are focused on testing the immunogenicity of these peptides in murine models, comparing those which contain overlapping and non-overlapping T and B cell epitopes. Results from such preclinical testing will inform an envisioned phase I clinical trial using a condensed peptide set targeting B cell epitopes with known viral neutralization plus optimal T cell epitopes.

Prior work has surveyed the epitope space of SARS-CoV-2 using analysis of sequence homology with SARS-CoV-1 epitopes, prediction of linear B cell epitopes, and prediction of T cell epitopes using IEDB tools. Grifoni *et al.* reported predicted T and B cell epitopes based on cross-referencing of known SARS epitopes with sequence homology to SARS-CoV-2 against SARS-CoV-2-specific parallel computational prediction^79^. However, this study did not consider epitope mapping of SARS-CoV-2 convalescent antibody repertoires, which may be important to achieve high specificity of B cell epitope predictions. Ahmed *et al.* reported a set of predicted T and B cell SARS-CoV-2 epitopes with associated assay confirmation within the NIAID ViPR database. However, these predicted epitopes were largely limited to those with sequence homology between SARS-CoV-1 and SARS-CoV-2, given the paucity of available SARS-CoV-2 assay data. Several studies have identified linear B cell epitopes on the SARS-CoV-2 surface glycoprotein from sera of viral exposed patients using peptide arrays^56–58^ as well as phage immunoprecipitation sequencing (PhIP-Seq)^59^. These studies are an important source of information but their results may include many epitopes from degraded proteins and thus would not be able to promote viral neutralization *in vivo* due to a lack of surface exposure. Our work adds to this important emerging field by analyzing the SARS-CoV-2 HLA ligand landscape through binding affinity filters derived from validated IEDB HLA ligands, as well as deriving T and B cell vaccine candidates through rational filtering criteria grounded in SARS-CoV-2 biology, including predicted immunogenicity, epitope location, glycosylation sites, and polymorphic sites. No other study to date has considered all such features in their epitope selection process. Additionally, inclusion of corresponding murine epitopes allows for future studies to be performed in animal models of SARS-CoV-2. We expect the application of these filters will improve specificity of antiviral response. As such, future studies testing the immunogenicity and efficacy of these filtered vaccine candidates in murine models will provide information critical in the design of a rationally optimized human SARS-CoV-2 vaccine.

Another unique aspect of our epitope selection process is the prioritization of overlapping CD4^+^, CD8^+^, and B cell epitopes. As the role of T cell epitope vaccines has not yet been clearly studied in SARS-CoV-2, we furthermore cross-referenced human and murine T cell epitopes to allow for murine vaccine studies using human-relevant peptides in H2-b and H2-d haplotypes. We hypothesize that inclusion of CD8^+^ epitopes may allow for clearance of SARS-CoV-2 from infected cells, and the inclusion of CD4^+^ epitopes may allow for greater activation of both cytotoxic and humoral antiviral responses. While overlapping CD4^+^ and CD8^+^ epitopes allowed for selection of peptide candidates covering a large proportion of the population, B/T cell overlapping epitope regions were more sparse due to the paucity of predicted B cell candidates. Thus, we expect the inclusion of overlapping CD4^+^/CD8^+^ optimized peptides alongside B cell optimized peptides to provide the most robust and broad antiviral adaptive immune coverage.

In addition to epitope selection, optional adjuvant choice for a SARS-CoV-2 vaccine is currently unclear. Current evidence suggests a Th2 dominant response to be associated with worse outcomes^9^ — thus, adjuvant selection may play an important role in skewing the helper arm toward a Th1 phenotype. Studies testing these questions are currently underway in murine models. Additionally, preliminary analysis of scRNA-seq dataset in ACE2 expressing cells of the respiratory^80–84^ and gastrointestinal^82,85,86^ tracts demonstrated increased expression of non-traditional checkpoint inhibitors (VISTA, Galectin 9, VTCN1), suggesting these as potential pathways to target for vaccine co-therapy (**Figure S10**). It remains unclear at this time if any of these above pathways are exploited by SARS-CoV-2 for innate or adaptive immune evasion.

One limitation of our study is that, while we use epitope mapping data with direct biological evidence for B cell epitopes in SARS-CoV-2, the T cell epitopes we report were all derived from computational prediction. In an effort to partially overcome this weakness, we applied binding affinity and immunogenicity prediction filters grounded in validated IEDB binding and tetramer studies. Reassuringly, the two extant studies examining T cell responses in COVID-19 patients have identified recurrent T cell epitopes which overlap with the vaccine peptides presented here. Le Bert *et al.* looked for T cell epitopes within the nucleocapsid (N), NSP-7 and NSP-13 proteins in PBMCs of recovered COVID-19 patients using an IFN-γ ELISpot assay^87^. They identified two recurrent epitope regions (N101-120, N321-340) which overlap with multiple 27mer vaccine peptides in this paper (**Figure 5B**, peptides 4-8). Shomuradova *et al.* also identified COVID-19 patient T cell epitopes, but using A*02:01 tetramers loaded with 13 distinct peptides from the surface glycoprotein (S)^88^. Two of these 13 peptides showed recurrent reactivity across 14 A*02:01 positive patients (S269-277 and S1000-1008). Both of these epitopes are also included in multiple 27mer vaccine peptides (Figure 5B, peptides 11 and 15).

Another potential limitation of this study is the insensitivity of our experiments to the total potential space of SARS-CoV-2 antibody epitopes. Our B cell epitope analyses start with only 58 identified linear antibody epitopes on the surface glycoprotein of SARS-CoV-2, while it is likely that many other epitopes are possible. Second, these linear epitope mappings do not allow for identification of antibodies which bind tertiary/quaternary protein structures. Lastly, identification of epitopes via array studies depended on differences in antibody binding to potential linear epitopes between uninfected and infected persons. There may be some cross-reactivity between antibodies generated against other coronaviruses and SARS-CoV-2, which if present might show reactivity in our screening assay. If true, our strategy would not identify these epitopes as specific for SARS-CoV-2. Similarly, we excluded viral regions with significant polymorphism across the viral population. As polymorphic regions may be under selection pressure, at least some of which may be due to antiviral immunity, these regions may prove to be better epitope targets in patients infected by the relevant viral strains. We have avoided these in the current study, however, as we have focused here on conserved regions of SARS-CoV-2 to identify epitopes that would be most broadly targetable in the human population. For these reasons, we do not present our antibody data as describing the complete set of SARS-CoV-2 epitopes.

A peptide vaccine targeting B cells, CD4^+^ T cells, and CD8^+^ T cells in parallel may prove an important part of a multifaceted response to the COVID-19 pandemic, as such an approach has a potentially favorable development timeline and the potential to avoid ADE by precisely directing the antibody response toward functional (neutralizing) regions. However, we emphasize that epitope selection is only one aspect of the problem, and a key question is whether a peptide vaccine can be sufficiently immunogenic. Adjuvant selection, conjugation to carriers such as KLH^89^ or rTTHC^90^, and prime/boost approaches using orthogonal platforms are all potential avenues to explore. We anticipate that the sets of vaccine peptides reported here may be valuable in the preclinical development of these approaches.

## Methods

### Antibody epitope curation

Linear B cell epitopes on the SARS-CoV-2 surface glycoprotein were curated from four published studies^56–59^. Three of these studies screened polyclonal sera of convalescent COVID-19 patients using either peptide arrays^56,58^ or phage immuno-precipation sequencing (PhIP-Seq)^59^. One study characterized the epitopes of monoclonal neutralizing antibodies^57^. Additionally, we were provided as-of-yet unpublished results from a study of sera from six SARS-CoV-2-naive patient sera and nine SARS-CoV-2-infected patient sera using PEPperCHIP® SARS-CoV-2 Proteome Microarrays. The peptides included in these proteome-wide epitope mapping analyses were limited to those which demonstrated either IgG or IgA fluorescence intensity > 1000U in at least two infected patient samples and in none of the naive patient samples. In addition, two peptides were also included (QGQTVTKKSAAEASK, QTVTKKSAAEASKKP) which demonstrated IgG fluorescence intensity > 1000U in only one naive patient sample each, but in four and five infected patient samples, respectively.

### HLA ligand prediction

The SARS-CoV-2 protein sequence FASTA was retrieved from the NCBI reference database (https://www.ncbi.nlm.nih.gov/nuccore/MT072688). Haplotypes included in this analysis were derived from those with > 5% expression within the United States populations based on the National Marrow Donor Program’s HaploStats tool^22^:

- **HLA-A**: A*11:01, A*02:01, A*01:01, A*03:01, A*24:02
- **HLA-B**: B*44:03, B*07:02, B*08:01, B*44:02, B*44:03, B*35:01
- **HLA-C**: C*03:04, C*04:01, C*05:01, C*06:02,C*07:01, C*07:02
- **HLA-DR**: DRB1*01:01, DRB1*03:01, DRB1*04:01, DRB1*07:01, DRB1*11:01, DRB1*13:01, DRB1*15:01

Additionally, HLA-DQ alpha/beta pairs were chosen based on prevalence in previous studies^23^:

- **HLA-DQ**: DQA1*01:02/DQB1*06:02, DQA1*05:01/DQB1*02:01, DQA1*02:01/DQB1*02:02, DQA1*05:05/DQB1*03:01, DQA1*01:01/DQB1*05:01, DQA1*03:01/DQB1*03:02, DQA1*03:03/DQB1*03:01, DQA1*01:03/DQB1*06:03

For HLA-I, 8-11mer epitopes were predicted using netMHCpan 4.0^18^ and MHCflurry 1.6.0^19^. For HLA-II caling, 15mers were predicted using NetMHCIIpan 3.2^20^ and NetMHCIIpan 4.0^21^. For optimization of epitope predictions, individual features from each HLA-I and HLA-II prediction tool was compared against IEDB binding affinities using Spearman correlation (**Figure S1)**. Cutpoints for the best performing HLA-I and HLA-II feature were set using 90% specificity of predicting for peptides with < 500nM binding affinity in the IEDB set. The proportion of the total U.S. population containing at least one haplotype capable of binding each peptide was calculated assuming no genetic linkage:

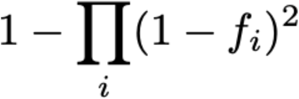

### Immunogenicity modeling

IEDB HLA-I and HLA-II viral tetramer data were used to generate a generalized linear model (GLM, family = binary) with tetramer-positivity as a binary outcome. Independent variables for HLA-I included NetMHCpan 4.0 binding affinity and elution score, MHCflurry binding affinity, presentation score, processing score, and percentage of aromatic (F, Y, W), acidic (D, E), basic (K, R H), small (A, G, S, T, P), cyclic (P), and thiol (C, M) amino acid residues. Independent variables for HLA-II included NetMHCIIpan 4.0 binding affinity and elution scores, and percentage of aromatic, acidic, basic, small, cyclic, and thiol amino acid residues. All independent variables were normalized to 0-1 to keep coefficients comparable (binding affinities divided by 50,000). GLM model performance was derived using 5-fold cross validation, balancing for HLA alleles. The final HLA-I and HLA-II models were generated using each full IEDB set, then applied to SARS-CoV-2 predicted HLA ligands to derive a GLM score. For immunogenicity filtering, predicted epitopes above the median GLM score were kept.

### SARS-CoV-2 entropy calculations

8,008 SARS-CoV-2 genome sequences were downloaded from GISAID (https://www.gisaid.org/)^51^. A preprocessing step removed 127 sequences that were shorter than 25,000 bases. The sequences were split into 79 smaller files and aligned using augur^52^ with MT072688.1^91^ as the reference genome. The reference genome was downloaded from NCBI GenBank^92^. The 79 resulting alignment files were concatenated into a single alignment file with the duplicate reference genome alignments removed. The multiple sequence alignment was translated to protein space using the R packages seqinr^93^ and msa^94^. Entropy for each position was calculated using the following formula, where *n* is the number of possible outcomes (i.e. total unique identifiable amino acid residues at each location) and *p*_*i*_ is the probability of each outcome (i.e. probability of each possible amino acid residues at each location):

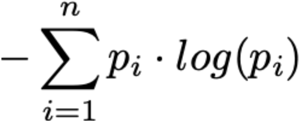

### Immunomodulatory molecule co-expression analysis

Single cell RNA sequencing data was collected from six respiratory datasets^80–84^ and three gastrointestinal datasets^82,85,86^. ACE2^+^ cells were subsetted as cells with an expression of ACE2 greater than zero. The proportion of ACE2^+^ cells expressing the immunomodulatory genes were plotted with the circlize package^95^. Coexpression of the immunomodulatory genes that were expressed in greater than five percent of the ACE2^+^ cells were plotted as links.

### Graphical and statistical analysis

Plots and analyses were generated using the following R packages: scales^96^, data.table^97^, ggrepel^98^, ggplot2^99^, viridis^100^, ggnewscale^101^, seqinr^93^, DESeq2^102^, GenomicRanges^103^, gplots^104^, ggbeeswarm^105^, ggallin^106^, stringr^107^, gridExtra^108^, pROC^109^, caret^110^, RColorBrewer^111^, dplyr^112^, cowplot^113^, ggpubr^114^, doMC^115^, venneuler^116^, ComplexHeatmap^117^, and circlize^95^ packages. Figures 4C, 4D, and 5 were generated using the following Python packages: NumPy^118^, pandas^119^, Matplotlib^120^, and Jupyter^121^.

## Code and Data availability

Data and analyses presented in this manuscript are available at: https://github.com/Benjamin-Vincent-Lab/Landscape-and-Selection-of-Vaccine-Epitopes-in-SARS-CoV-2

Several data files larger than 100Mb and supplemental tables are available at: https://data.mendeley.com/datasets/c6pdfrwxgj/2

## Acknowledgements

The authors appreciate funding support from University of North Carolina University Cancer Research Fund (AR and BGV), the Susan G. Komen Foundation (BGV), the V Foundation for Cancer Research (BGV), and the National Institutes of Health (CCS, 1F30CA225136). We would like to thank members of the #DownWithTheCrown Slack channel for helpful discussion and feedback.

