## Supplemental Figures S1 - S10 for "Landscape and Selection of Vaccine Epitopes in SARS-CoV-2"

### Supplemental materials

Supplemental tables S1-S5 are available at: https://data.mendeley.com/datasets/c6pdfrwxgj/2

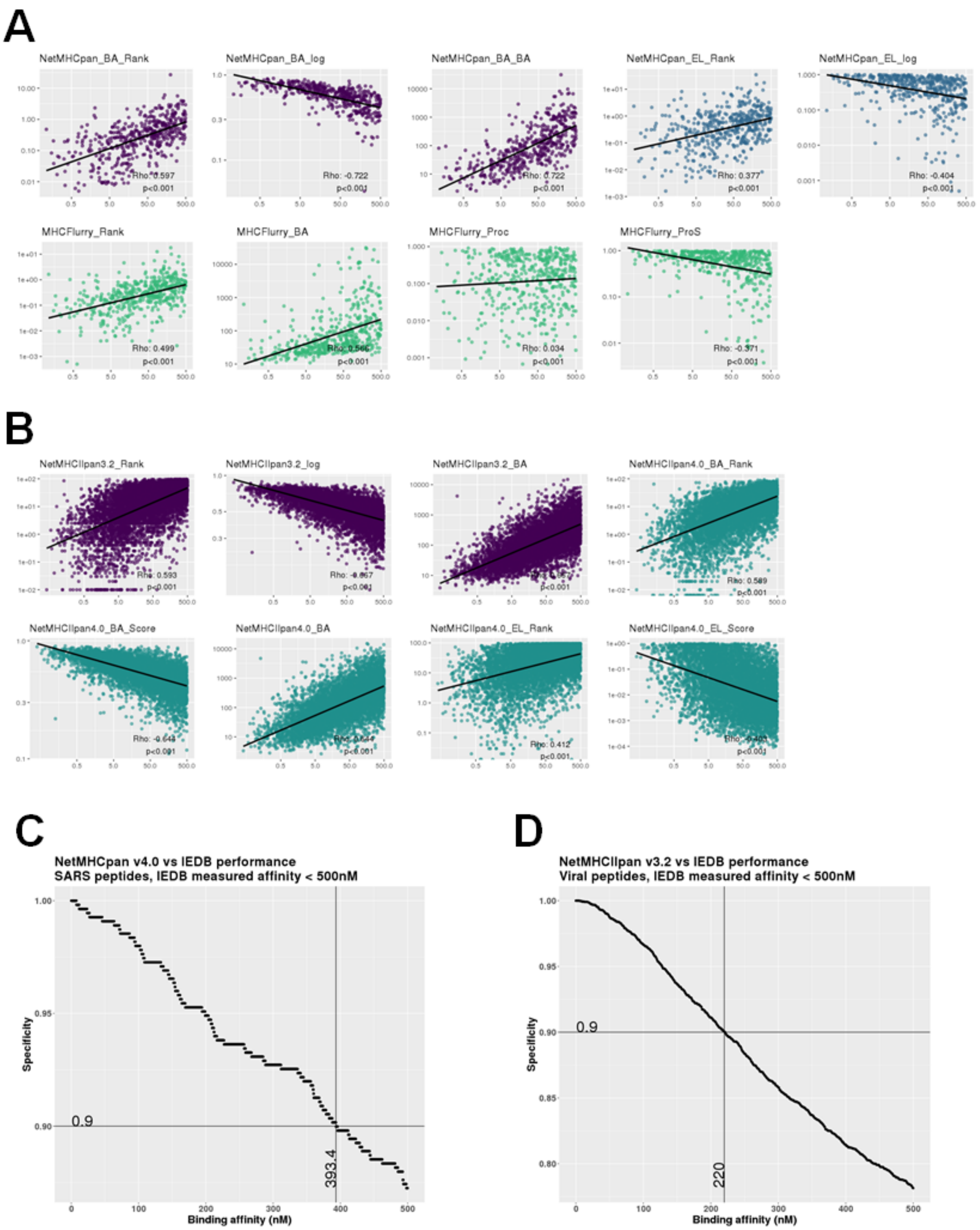


***Figure S1: Selection criteria for predicted HLA ligands.*** ***(A&B)*** *Scatterplot of IEDB binding affinity (x-axis) versus predicted features (y-axis) for HLA-I SARS ligands* ***(A)*** *and HLA-II viral ligands* ***(B)****, with linear fit and Spearman correlation represented. Color represents the prediction tool used for each feature.* ***(C&D)*** *Plot of NetMHCpan 4.0* ***(C)*** *and NetMHCIIpan 3.2* ***(D)*** *binding affinity (x-axis) versus specificity (y-axis) for predicting binding ligand, as defined by IEDB binding affinity < 500 nM.*

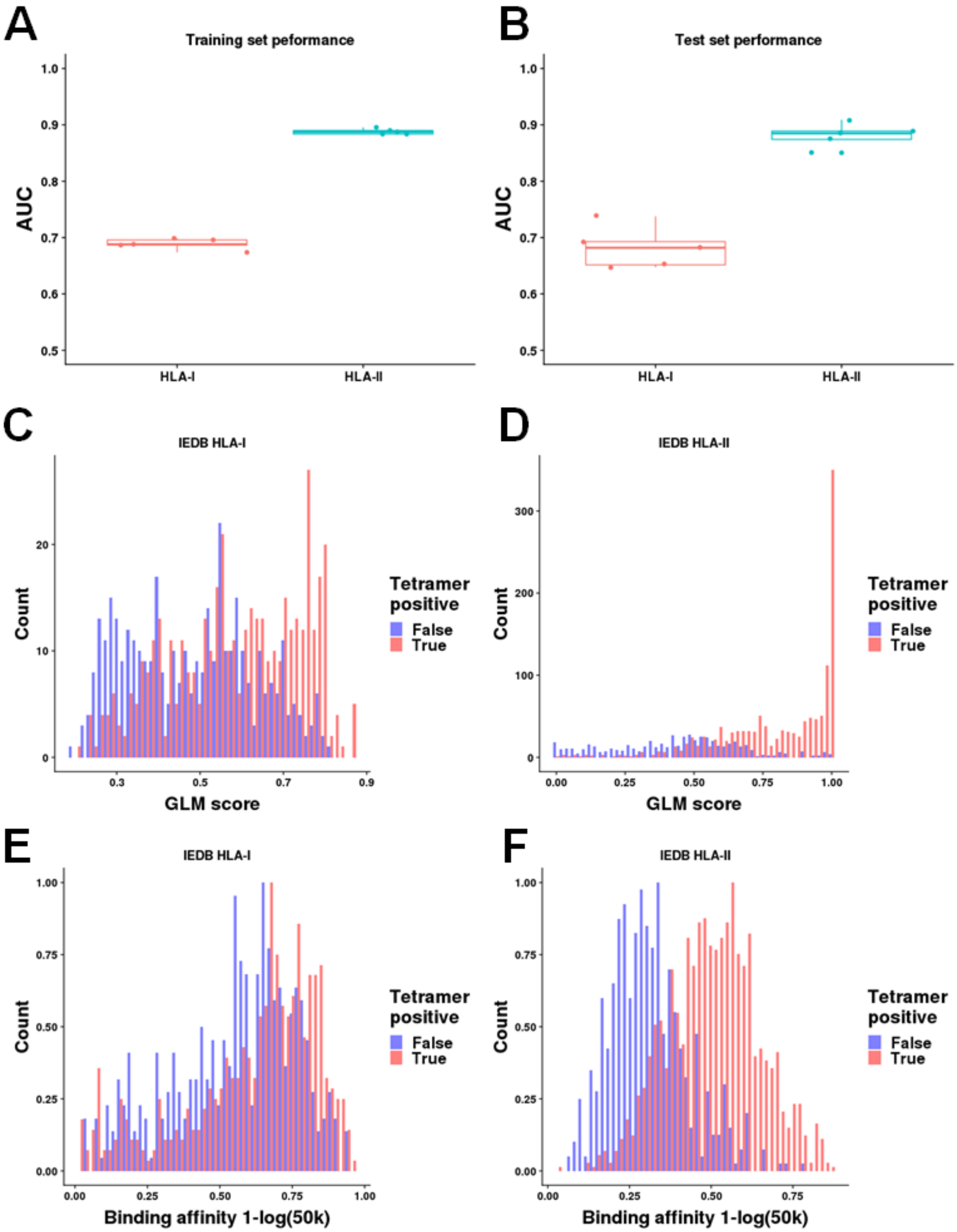


***Figure S2: Summary of multivariable GLM model for prediction of epitope immunogenicity, trained on IEDB tetramer data.*** ***(A&B)*** *Area under the curve of HLA-I (red) and HLA-II (blue) GLM models for 5-fold cross validation training* ***(A)*** *and test* ***(B)*** *sets.* ***(C&D)*** *Histograms of GLM scores for tetramer positive (red) and negative (blue) CD8$^+$* ***(C)*** *and CD4$^+$* ***(D)*** *epitopes in IEDB tetramer dataset.* ***(E&F)*** *Histograms of binding affinity scores for tetramer positive (red) and negative (blue) CD8$^+$* ***(E)*** *and CD4$^+$* ***(F)*** *epitopes in IEDB tetramer dataset.*

A

```
Deviance Residuals:
    Min       1Q   Median       3Q      Max
-1.8262  -1.0980   0.6825   1.0449   1.7833

Coefficients:
                  Estimate Std. Error z value Pr(>|z|)
(Intercept)        -0.7444     0.2108  -3.531 0.000413 ***
Flurry_proc_score   1.0901     0.3146   3.465 0.000530 ***
EL_Score            1.4007     0.2876   4.870 1.11e-06 ***
Binding_affinity    3.1258     0.8026   3.894 9.84e-05 ***
Small              -1.1527     0.4761  -2.421 0.015457 *
---
Signif. codes:  0 '***' 0.001 '**' 0.01 '*' 0.05 '.' 0.1 ' ' 1

(Dispersion parameter for binomial family taken to be 1)

    Null deviance: 1117.2  on 808  degrees of freedom
Residual deviance: 1025.9  on 804  degrees of freedom
AIC: 1035.9

Number of Fisher Scoring iterations: 4
```

B

```
Deviance Residuals:
    Min       1Q   Median       3Q      Max
-4.1923  -0.6214   0.1554   0.7503   5.0196

Coefficients:
                 Estimate Std. Error z value Pr(>|z|)
(Intercept)        1.4417     0.2348   6.139 8.30e-10 ***
EL_Score           9.5627     0.9015  10.608  < 2e-16 ***
Binding_affinity -17.5174     2.0503  -8.544  < 2e-16 ***
Cyclic            -5.2677     1.2746  -4.133 3.59e-05 ***
Aromatic          -2.7801     0.9348  -2.974  0.00294 **
Acidic            -2.2112     0.8657  -2.554  0.01065 *
Basic             -1.4255     0.7661  -1.861  0.06278 .
---
Signif. codes:  0 '***' 0.001 '**' 0.01 '*' 0.05 '.' 0.1 ' ' 1

(Dispersion parameter for binomial family taken to be 1)

    Null deviance: 2285.8  on 1859  degrees of freedom
Residual deviance: 1504.4  on 1853  degrees of freedom
AIC: 1518.4
```

***Figure S3: (A&B)*** *HLA-I* ***(A)*** *and HLA-II* ***(B)*** *GLM predicting for tetramer positivity as a function of binding and amino acid features.*

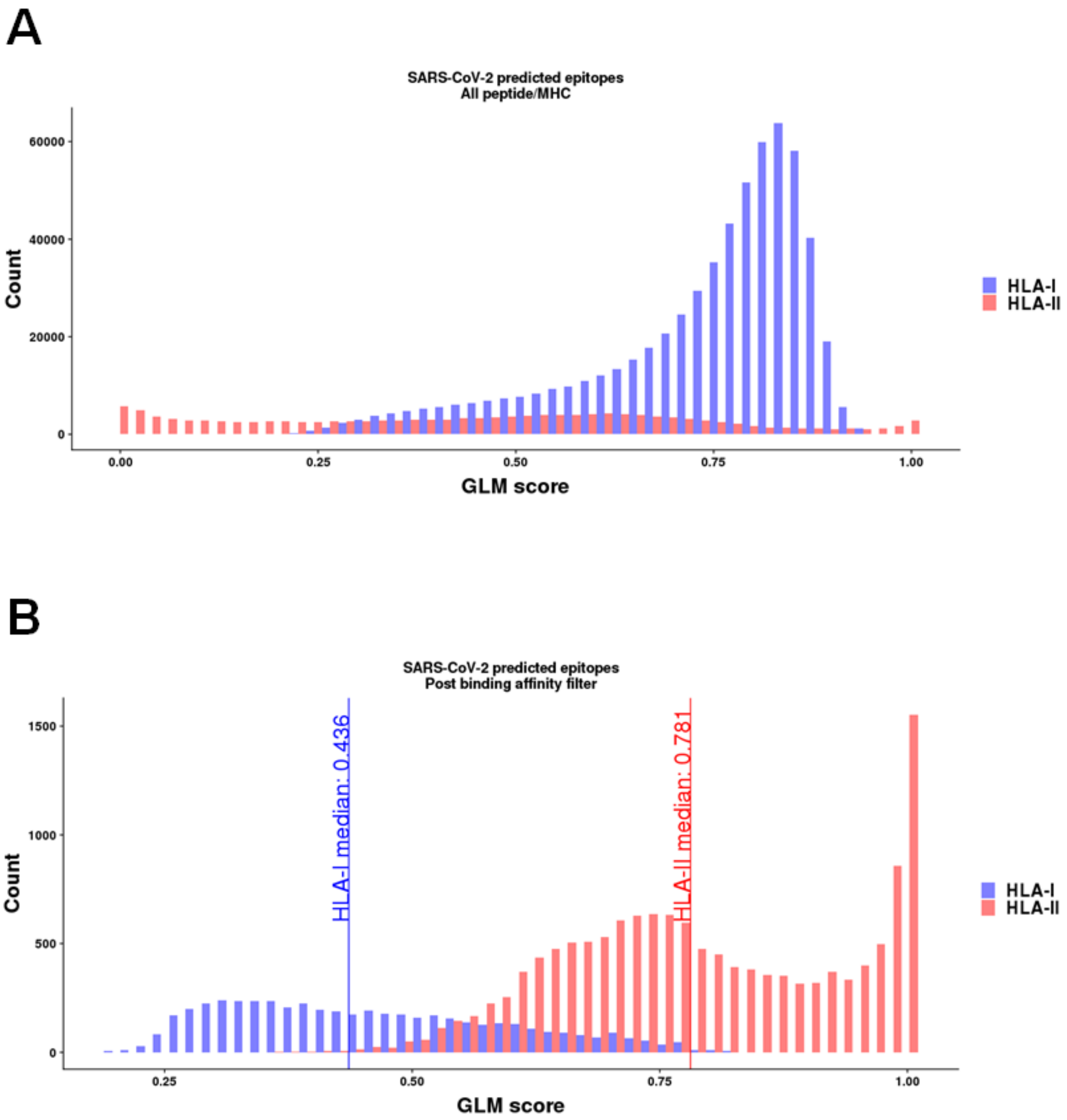


***Figure S4: (A&B)*** *Distribution of GLM scores among predicted SARS-CoV-2 T cell epitopes prior to binding affinity filter* ***(A)*** *and after binding affinity filter* ***(B)****. Vertical lines in* ***(B)*** *represent median GLM score for predicted $CD4^+$ and $CD8^+$ epitopes.*

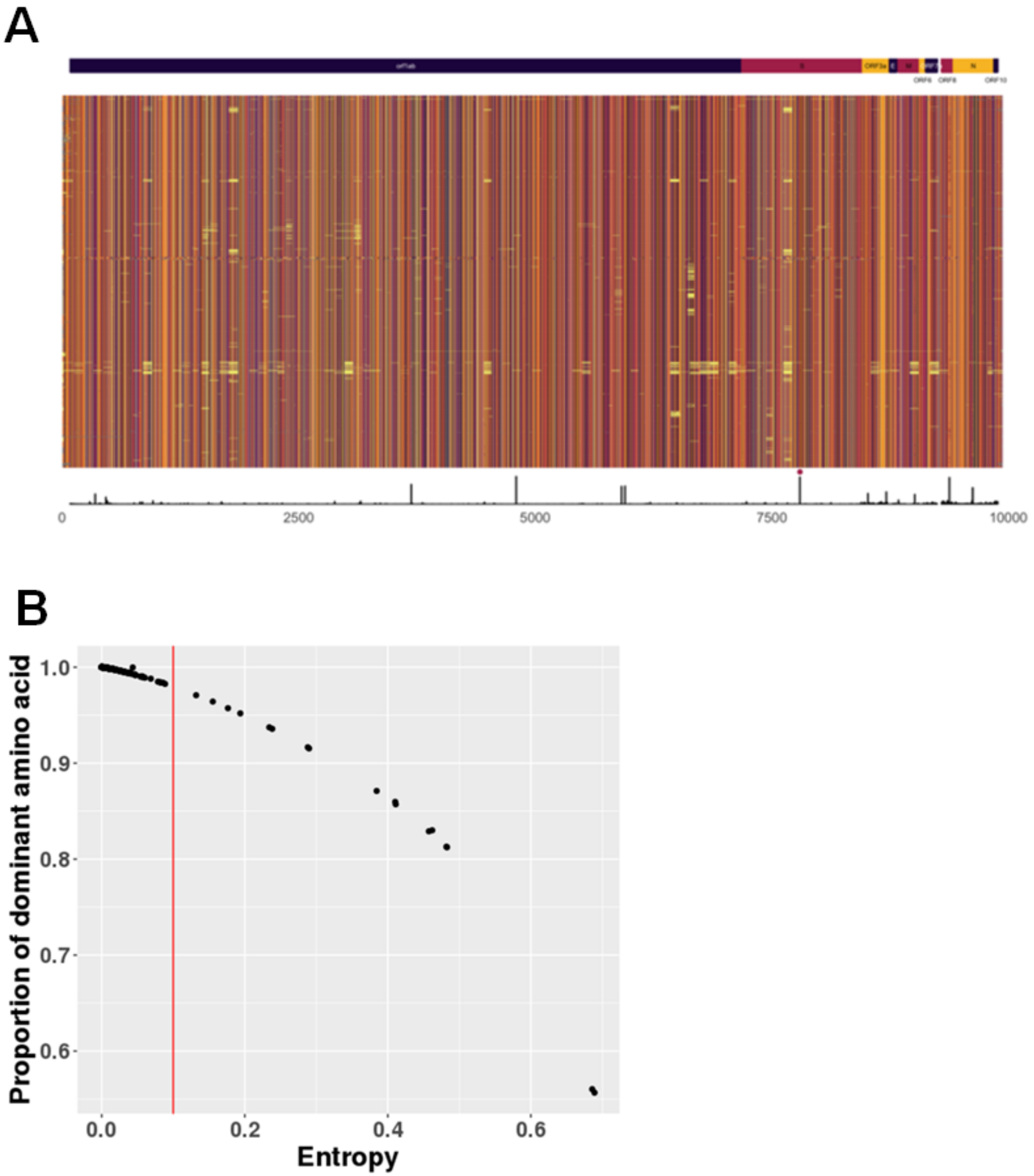


***Figure S5: Sequence level variation across SARS-CoV-2 viral proteomes in the Nextstrain database. (A)*** *Locations along the viral genome represented by x-axis, with individual genomes (n=7882) along y-axis. Colors represent amino acid residues (plotted on viridis "inferno" color scheme, dark (A) to light (Y) in alphabetical order of amino acid letter abbreviations; gap/unknown = grey), aligned using multiple sequence alignment (MSA). Histogram along the y-axis represents entropy at each location, with position 614 of S protein marked with red dot. Proteins by locations are shown by column-side colorbar.* ***(B)*** *Entropy versus proportion of dominant amino acid residue by position along MSA-aligned genomes, with red line representing entropy cutoff of 0.1 used for this study.*

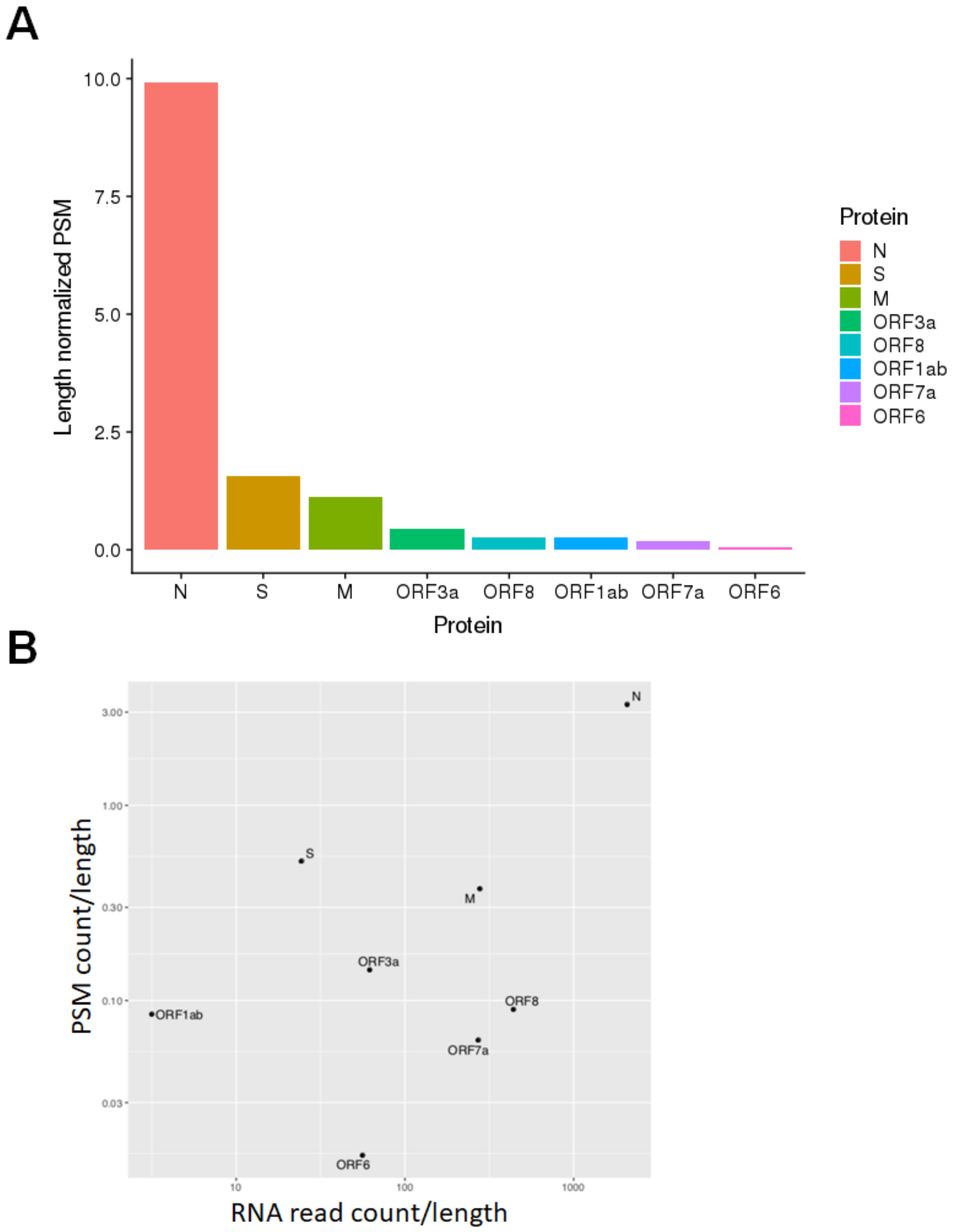


***Figure S6: (A)*** *Length normalized peptide spectrum match (PSM) counts for SARS-CoV-2 proteins.* ***(B)*** *Length normalized PSM versus length normalized RNA-seq read counts for SARS-CoV-2 proteins.*

**A**

| **Peptide Feature** | **Difficulty Score** |
|---|---|
| Entire peptide hydrophobic (GRAVY score > 2.0) | 1 |
| Difficult N-terminal residue | 1 |
| Difficult C-terminal residue | 2 |
| Number of cysteine or methionine residues | 2 |
| Difficult local hydrophobicity (local GRAVY score > 1.5) | 2 |
| Moderately unstable di-peptides | 3 |
| Disulfide bonds (more than one cysteine) | 5 |
| Extreme local hydrophobicity | 10 |
| Extremely unstable di-peptides | 10 |

**B**

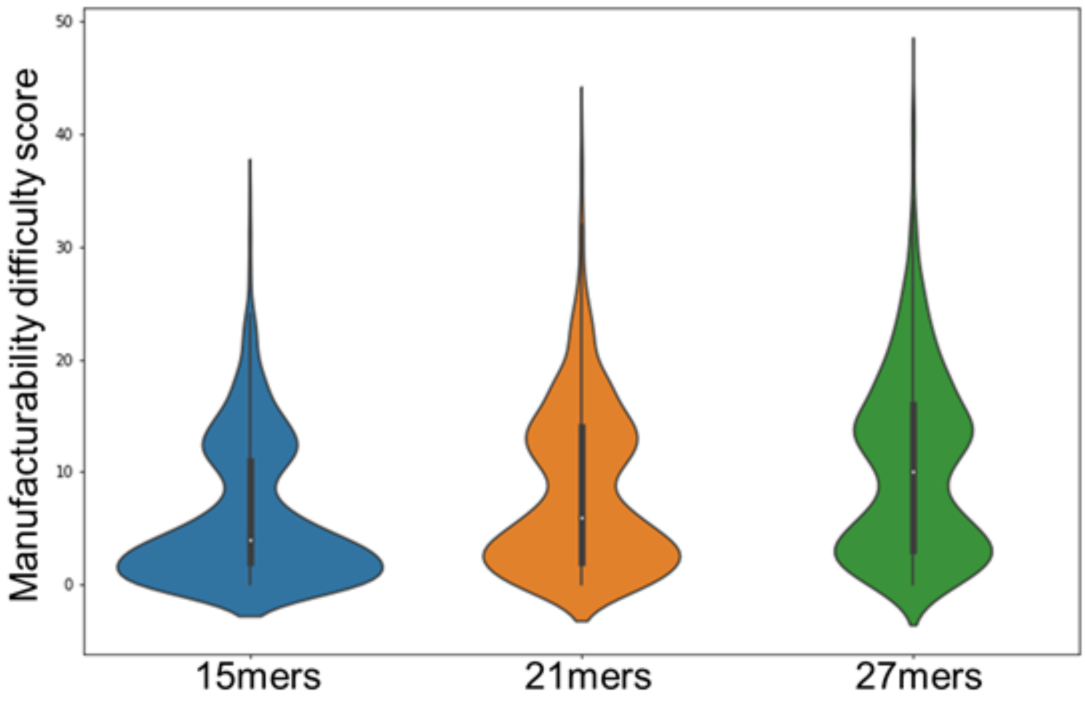


***Figure S7: (A)*** *Manufacturability difficulty scoring criteria for vaccine peptide candidates.* ***(B)*** *Distribution of manufacturability difficulty scores for 15mer, 21mer, and 27mer peptide sets.*

# A

| Symbol | Set | # Peptides | HLA-I Coverage | HLA-II Coverage | Total Coverage | # B-cell Epitope Regions |
|---|---|---|---|---|---|---|
| ⊛ | CD4+/CD8+ | 3 | 84.9% | 84.7% | 71.9% | 0 |
| ⊛$^{d}$ | CD4+/CD8+ (H2$^{d}$ ligands) | 4 | 90.2% | 84.7% | 76.4% | 0 |
| ⊛$^{b}$ | CD4+/CD8+ (H2$^{b}$ ligands) | 4 | 93.9% | 84.7% | 79.5% | 0 |
| ⊛$^{bd}$ | CD4+/CD8+ (H2$^{b}$ and H2$^{d}$ ligands) | 4 | 92.1% | 84.7% | 78.0% | 0 |
| ○ | CD4+ | 3 | 92.2% | 88.5% | 81.6% | 0 |
| ○$^{d}$ | CD4+ (H2$^{d}$ ligands) | 3 | 92.2% | 88.5% | 81.6% | 0 |
| ○$^{b}$ | CD4+ (H2$^{b}$ ligands) | 3 | 69.5% | 84.7% | 58.9% | 0 |
| ○$^{bd}$ | CD4+ (H2$^{b}$ and H2$^{d}$ ligands) | 3 | 93.8% | 84.7% | 79.4% | 0 |
| * | CD8+ | 3 | 95.1% | 62.2% | 59.1% | 0 |
| *$^{d}$ | CD8+ (H2$^{d}$ ligands) | 3 | 94.7% | 68.9% | 65.3% | 0 |
| *$^{b}$ | CD8+ (H2$^{b}$ ligands) | 3 | 94.7% | 68.9% | 65.3% | 0 |
| *$^{bd}$ | CD8+ (H2$^{b}$ and H2$^{d}$ ligands) | 3 | 94.7% | 68.9% | 65.3% | 0 |
| ⧈ | B-Cell/CD4+/CD8+ | 4 | 84.2% | 62.7% | 52.8% | 3 |
| ⊡ | B-Cell/CD4+ | 4 | 84.2% | 62.7% | 52.8% | 3 |
| ⊡$^{b}$ | B-Cell/CD4+ (H2$^{b}$ ligands) | 2 | 37.2% | 39.4% | 14.6% | 2 |
| ⧆ | B-Cell/CD8+ | 3 | 88.9% | 22.6% | 20.1% | 3 |
| ⧆$^{d}$ | B-Cell/CD8+ (H2$^{d}$ ligands) | 1 | 77.2% | 38.4% | 29.7% | 1 |
| ⧆$^{b}$ | B-Cell/CD8+ (H2$^{b}$ ligands) | 2 | 87.2% | 22.6% | 19.7% | 2 |
| ⧆$^{bd}$ | B-Cell/CD8+ (H2$^{b}$ and H2$^{d}$ ligands) | 1 | 77.2% | 38.4% | 29.7% | 1 |
| □ | B-Cell | 3 | 78.0% | 40.7% | 31.8% | 3 |

# B

| | Sequence | Protein | Start | End | B-cell Epitope Region | HLA-I Coverage | HLA-II Coverage | H2$^{b}$ I | H2$^{b}$ II | H2$^{d}$ I | H2$^{d}$ II | Selection Sets |
|---|---|---|---|---|---|---|---|---|---|---|---|---|
| 1 | LLQFAYANRNRFLYIIKLIFL | M | 34 | 54 | | 89.0% | 36.0% | + | + | + | + | * *$^{b}$ *$^{d}$ *$^{bd}$ ○ ○$^{d}$ ○$^{bd}$ ⊛$^{d}$ ⊛$^{b}$ ⊛$^{bd}$ |
| 2 | FVLAAVYRINWITGGIAIAMA | M | 65 | 85 | | 42.0% | 76.0% | + | + | - | + | ○$^{b}$ ⊛ ⊛$^{b}$ |
| 3 | LSYFIASFRLFARTRSMWSFN | M | 93 | 113 | | 78.0% | 46.0% | + | + | + | + | ⊛$^{bd}$ |
| 4 | LSPRWYFYYLGTGPEAGLPYG | N | 104 | 124 | | 49.0% | 23.0% | + | + | + | - | * |
| 5 | GTRNPANNAAIVLQLPQGTTL | N | 147 | 167 | | 20.0% | 55.0% | - | + | - | + | ○$^{bd}$ |
| 6 | IAQFAPSASAFFGMSRIGMEV | N | 304 | 324 | | 63.0% | 51.0% | + | + | + | + | ⊛$^{d}$ ⊛$^{bd}$ |
| 7 | SASAFFGMSRIGMEVTPSGTW | N | 310 | 330 | | 65.0% | 37.0% | + | - | + | - | *$^{b}$ *$^{d}$ *$^{bd}$ |
| 8 | IGMEVTPSGTWLTYTGAIKLD | N | 320 | 340 | | 54.0% | 52.0% | + | + | - | - | ⊛$^{b}$ |
| 9 | GTWLTYTGAIKLDDKDPNFKD | N | 328 | 348 | | 26.0% | 62.0% | + | + | - | - | ○$^{b}$ ⊛ |
| 10 | KQQTVTLLPAADLDDFSKQLQ | N | 388 | 408 | | 11.0% | 52.0% | - | - | - | + | ○ ○$^{d}$ |
| 11 | LPFNDGVYFASTEKSNIIRGW | S | 84 | 104 | | 58.0% | 41.0% | - | + | - | - | * |
| 12 | PLVDLPIGINITRFQTLLALH | S | 225 | 245 | | 65.0% | 62.0% | + | - | + | + | ⊛ ⊛$^{d}$ |
| 13 | GAAAYYVGYLQPRTFLLKYNE | S | 261 | 281 | | 88.0% | 38.0% | + | + | + | - | *$^{b}$ *$^{d}$ *$^{bd}$ |
| 14 | LTDEMIAQYTSALLAGTITSG | S | 865 | 885 | | 42.0% | 73.0% | + | + | + | + | ⊛$^{d}$ ⊛$^{bd}$ |
| 15 | LSSNFGAISSVLNDILSRLDK | S | 966 | 986 | | 59.0% | 62.0% | + | + | - | + | ⊛$^{b}$ |
| 16 | VTQQLIRAAEIRASANLAATK | S | 1008 | 1028 | | 30.0% | 81.0% | - | + | - | + | ○ ○$^{d}$ ○$^{b}$ ○$^{bd}$ |
| 17 | NYNYLYRLFRKSNLKPFERDI | S | 448 | 468 | 456-FRKSNLKPFERDISTEIY-473 | 77.0% | 38.0% | + | - | + | - | ⧆$^{d}$ ⧆$^{bd}$ ⊡ ⧈ |
| 18 | YRLFRKSNLKPFERDISTEIY | S | 453 | 473 | 456-FRKSNLKPFERDISTEIY-473 | 78.0% | 23.0% | + | - | - | - | □ ⧆ ⧆$^{b}$ ⊡ ⧈ |
| 19 | KPFERDISTEIYQAGSTPCNG | S | 462 | 482 | 456-FRKSNLKPFERDISTEIY-473 | 20.0% | 21.0% | - | + | - | - | ⊡$^{b}$ |
| 20 | QFGRDIADTTDAVRDPQTLEI | S | 564 | 584 | 580-QTLE-583 | 0.0% | 0.0% | - | - | - | - | □ |
| 21 | PQTLEILDITPCSFGGVSVIT | S | 579 | 599 | 580-QTLE-583 | 13.0% | 0.0% | - | - | - | - | ⧆ |
| 22 | QTLEILDITPCSFGGVSVITP | S | 580 | 600 | 580-QTLE-583 | 13.0% | 21.0% | - | - | - | - | ⊡ ⧈ |
| 23 | GFNFSQILPDPSKPSKRSFIE | S | 799 | 819 | 809-PSKP-812 | 21.0% | 23.0% | - | + | - | - | □ ⊡ ⊡$^{b}$ ⧈ |
| 24 | PSKPSKRSFIEDLLFNKVTLA | S | 809 | 829 | 809-PSKP-812 | 66.0% | 0.0% | + | - | - | - | ⧆ ⧆$^{b}$ |

***Figure S8: T cell and B cell vaccine candidates.*** ***(A)*** *21mer vaccine peptide sets selecting for best CD4$^{+}$, CD8$^{+}$, CD4$^{+}$/CD8$^{+}$, and B cell epitopes with HLA-I, HLA-II, and total population coverage.* ***(B)*** *Unified list of all selected 21mer vaccine peptides. Vaccine peptides containing predicted ligands for murine MHC alleles (H2-b and H2-d haplotypes) are indicated in their respective columns.*

## A

| Symbol | Set | # Peptides | HLA-I Coverage | HLA-II Coverage | Total Coverage | # B-cell Epitope Regions |
|---|---|---|---|---|---|---|
| ⊛ | CD4+/CD8+ | 5 | 90.6% | 88.5% | 80.2% | 0 |
| ⊛$^{d}$ | CD4+/CD8+ (H2$^{d}$ ligands) | 3 | 81.1% | 76.2% | 61.8% | 0 |
| ⊛$^{b}$ | CD4+/CD8+ (H2$^{b}$ ligands) | 3 | 81.8% | 62.5% | 51.1% | 0 |
| ⊛$^{bd}$ | CD4+/CD8+ (H2$^{b}$ and H2$^{d}$ ligands) | 2 | 77.2% | 65.8% | 50.8% | 0 |
| ○ | CD4+ | 3 | 83.9% | 88.5% | 74.3% | 0 |
| ○$^{d}$ | CD4+ (H2$^{d}$ ligands) | 3 | 86.7% | 84.7% | 73.4% | 0 |
| ○$^{b}$ | CD4+ (H2$^{b}$ ligands) | 3 | 83.9% | 84.7% | 71.1% | 0 |
| ○$^{bd}$ | CD4+ (H2$^{b}$ and H2$^{d}$ ligands) | 3 | 86.7% | 84.7% | 73.4% | 0 |
| ∗ | CD8+ | 3 | 95.8% | 38.4% | 36.8% | 0 |
| ∗$^{d}$ | CD8+ (H2$^{d}$ ligands) | 3 | 94.6% | 22.6% | 21.4% | 0 |
| ∗$^{b}$ | CD8+ (H2$^{b}$ ligands) | 3 | 91.2% | 46.5% | 42.4% | 0 |
| ∗$^{bd}$ | CD8+ (H2$^{b}$ and H2$^{d}$ ligands) | 3 | 91.2% | 46.5% | 42.4% | 0 |
| [⊛] | B-Cell/CD4+/CD8+ | 4 | 77.2% | 45.8% | 35.3% | 3 |
| ⧇ | B-Cell/CD4+ | 5 | 77.2% | 62.7% | 48.4% | 3 |
| ⧇$^{b}$ | B-Cell/CD4+ (H2$^{b}$ ligands) | 2 | 0.0% | 39.4% | 0.0% | 2 |
| ⧆ | B-Cell/CD8+ | 6 | 84.2% | 29.9% | 25.2% | 3 |
| ⧆$^{d}$ | B-Cell/CD8+ (H2$^{d}$ ligands) | 1 | 77.2% | 20.4% | 15.8% | 1 |
| ⧆$^{b}$ | B-Cell/CD8+ (H2$^{b}$ ligands) | 2 | 72.5% | 20.4% | 14.8% | 1 |
| ⧆$^{bd}$ | B-Cell/CD8+ (H2$^{b}$ and H2$^{d}$ ligands) | 1 | 77.2% | 20.4% | 15.8% | 1 |
| □ | B-Cell | 3 | 44.0% | 11.9% | 5.2% | 3 |

## B

| | Sequence | Protein | Start | End | B-cell Epitope Region | HLA-I Coverage | HLA-II Coverage | H2$^{b}$ I | H2$^{b}$ II | H2$^{d}$ I | H2$^{d}$ II | Selection Sets |
|---|---|---|---|---|---|---|---|---|---|---|---|---|
| 1 | LLQFAYANRNRFLYI | M | 34 | 48 | | 77.0% | 36.0% | + | + | + | + | ○ ○$^{b}$ ○$^{d}$ ○$^{bd}$ ⊛ ⊛$^{d}$ ⊛$^{bd}$ |
| 2 | YANRNRFLYIIKLIF | M | 39 | 53 | | 78.0% | 0.0% | + | - | + | - | ∗$^{d}$ |
| 3 | ANRNRFLYIIKLIFL | M | 40 | 54 | | 81.0% | 0.0% | + | - | + | - | ∗$^{b}$ ∗$^{bd}$ |
| 4 | YFIASFRLFARTRSM | M | 95 | 109 | | 78.0% | 20.0% | + | - | + | + | ∗ |
| 5 | SFRLFARTRSMWSFN | M | 99 | 113 | | 73.0% | 46.0% | + | + | - | + | ⊛$^{b}$ |
| 6 | LSPRWYFYYLGTGPE | N | 104 | 118 | | 49.0% | 0.0% | + | - | + | - | ∗$^{d}$ ∗$^{b}$ ∗$^{bd}$ |
| 7 | ATKAYNVTQAFGRRG | N | 264 | 278 | | 24.0% | 46.0% | + | + | + | - | ⊛$^{b}$ |
| 8 | PQIAQFAPSASAFFG | N | 302 | 316 | | 17.0% | 39.0% | - | + | + | + | ○$^{d}$ ○$^{bd}$ ⊛$^{d}$ |
| 9 | SASAFFGMSRIGMEV | N | 310 | 324 | | 56.0% | 37.0% | + | - | + | - | ⊛ |
| 10 | MEVTPSGTWLTYTGA | N | 322 | 336 | | 46.0% | 0.0% | - | - | - | - | ∗ |
| 11 | PSGTWLTYTGAIKLD | N | 326 | 340 | | 14.0% | 52.0% | + | + | - | - | ○$^{b}$ |
| 12 | QQTVTLLPAADLDDF | N | 389 | 403 | | 11.0% | 34.0% | - | - | - | - | ○⊛ |
| 13 | IGINITRFQTLLALH | S | 231 | 245 | | 61.0% | 62.0% | + | - | + | + | ⊛⊛$^{d}$ |
| 14 | YYVGYLQPRTFLLKY | S | 265 | 279 | | 88.0% | 23.0% | - | + | + | - | ∗∗$^{d}$ |
| 15 | LTDEMIAQYTSALLA | S | 865 | 879 | | 42.0% | 46.0% | + | + | + | + | ∗$^{b}$ ∗$^{bd}$ ⊛$^{b}$ ⊛$^{bd}$ |
| 16 | RAAEIRASANLAATK | S | 1014 | 1028 | | 30.0% | 79.0% | - | + | - | + | ○ ○$^{b}$ ○$^{d}$ ○$^{bd}$ ⊛ |
| 17 | GGNYNYLYRLFRKSN | S | 446 | 460 | 456-FRKSNLKPFERDISTEIY-473 | 37.0% | 20.0% | + | - | + | - | ⧆ |
| 18 | NYNYLYRLFRKSNLK | S | 448 | 462 | 456-FRKSNLKPFERDISTEIY-473 | 77.0% | 20.0% | + | - | + | - | ⧆$^{d}$ ⧆$^{bd}$ ⧇[⊛] |
| 19 | YNYLYRLFRKSNLKP | S | 449 | 463 | 456-FRKSNLKPFERDISTEIY-473 | 73.0% | 20.0% | + | - | - | - | ⧆$^{b}$ |
| 20 | YLYRLFRKSNLKPFE | S | 451 | 465 | 456-FRKSNLKPFERDISTEIY-473 | 73.0% | 20.0% | + | - | - | - | ⧆ |
| 21 | YRLFRKSNLKPFERD | S | 453 | 467 | 456-FRKSNLKPFERDISTEIY-473 | 73.0% | 23.0% | + | - | - | - | ⧇[⊛] |
| 22 | RLFRKSNLKPFERDI | S | 454 | 468 | 456-FRKSNLKPFERDISTEIY-473 | 56.0% | 0.0% | + | - | - | - | ⧆$^{b}$ |
| 23 | FRKSNLKPFERDIST | S | 456 | 470 | 456-FRKSNLKPFERDISTEIY-473 | 32.0% | 0.0% | - | - | - | - | ⧆ |
| 24 | KSNLKPFERDISTEI | S | 458 | 472 | 456-FRKSNLKPFERDISTEIY-473 | 29.0% | 0.0% | - | - | - | - | □ |
| 25 | LKPFERDISTEIYQA | S | 461 | 475 | 456-FRKSNLKPFERDISTEIY-473 | 20.0% | 12.0% | - | - | - | - | ⧆ |
| 26 | ISTEIYQAGSTPCNG | S | 468 | 482 | 456-FRKSNLKPFERDISTEIY-473 | 0.0% | 21.0% | - | + | - | - | ⧇⧇$^{b}$ |
| 27 | ADTTDAVRDPQTLEI | S | 570 | 584 | 580-QTLE-583 | 0.0% | 0.0% | - | - | - | - | □ ⧇ [⊛] |
| 28 | PQTLEILDITPCSFG | S | 579 | 593 | 580-QTLE-583 | 13.0% | 0.0% | - | - | - | - | ⧆ |
| 29 | GFNFSQILPDPSKPS | S | 799 | 813 | 809-PSKP-812 | 0.0% | 23.0% | - | + | - | - | ⧇⧇$^{b}$ |
| 30 | FNFSQILPDPSKPSK | S | 800 | 814 | 809-PSKP-812 | 21.0% | 12.0% | - | - | - | - | □ ⧆ [⊛] |

***Figure S9: T cell and B cell vaccine candidates.*** ***(A)*** *15mer vaccine peptide sets selecting for best CD4$^{+}$, CD8$^{+}$, CD4$^{+}$/CD8$^{+}$, and B cell epitopes with HLA-I, HLA-II, and total population coverage.* ***(B)*** *Unified list of all selected 15mer vaccine peptides. Vaccine peptides containing predicted ligands for murine MHC alleles (H2-b and H2-d haplotypes) are indicated in their respective columns.*

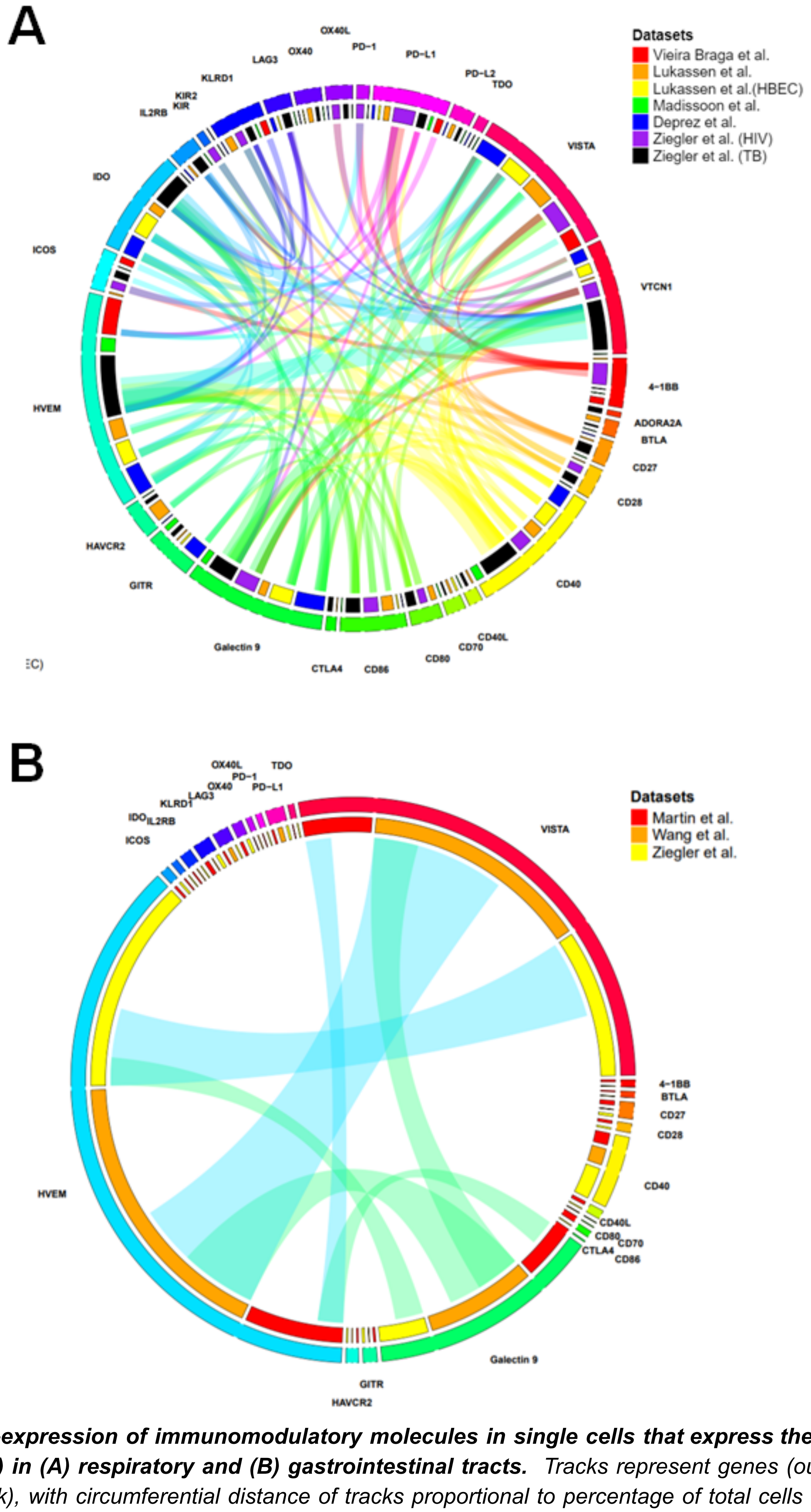


***Figure S10: Co-expression of immunomodulatory molecules in single cells that express the SARS-CoV-2 receptor (ACE2) in (A) respiratory and (B) gastrointestinal tracts.*** *Tracks represent genes (outer track) and study (inner track), with circumferential distance of tracks proportional to percentage of total cells which express each respective gene. Inner arcs represent co-expression of two genes, with width or arcs proportional to cell number. Arcs are filtered by co-expression >5% of ACE2 positive cells per dataset.*
